# Genomic and Phenomic Prediction for Soybean Seed Yield, Protein, and Oil

**DOI:** 10.1101/2024.11.01.621550

**Authors:** Liza Van der Laan, Kyle Parmley, Mojdeh Saadati, Hernan Torres Pacin, Srikanth Panthulugiri, Soumik Sarkar, Baskar Ganapathysubramanian, Aaron Lorenz, Asheesh K. Singh

## Abstract

Developments in genomics and phenomics have provided valuable tools for use in cultivar development. Genomic prediction (GP) has been used in commercial soybean [*Glycine max* L. (Merr.)] breeding programs to predict grain yield and seed composition traits. Phenomic prediction (PP) is a rapidly developing field that holds the potential to be used for the selection of genotypes early in the growing season. The objectives of this study were to compare the use and performance of GP and PP for predicting soybean seed yield, protein content, and oil content. We additionally conducted Genome Wide Association Studies (GWAS) to identify significant SNPs associated with the traits of interest. These SNPs were also used to train the GP models. The GWAS panel of 292 diverse accessions was grown in six environments in replicated trials. Spectral data were collected at three timepoints during the growing season. A GBLUP model was trained on 268 accessions, while three separate machine learning (ML) models were trained on vegetation indices (VIs) and canopy traits. We observed that for PP, Random Forest (RF) algorithm had the highest rank correlation between the predicted and the actual phenotype rank. PP had a higher correlation coefficient than GP for seed yield, while GP had higher correlation coefficients for seed protein and oil contents. VIs with high feature importance were used as covariates in a new GBLUP model, and a new RF model was trained with the inclusion of selected SNPs from the GWAS results. These models did not outperform the original GP and PP models. These results show the capability of using ML for in-season predictions for specific traits in soybean breeding and provide insights on PP and GP inclusions in breeding programs.

**PLAIN LANGUAGE SUMMARY:** Obtaining DNA information can often be costly for a public breeding program to obtain, which is a barrier of entry for using genomic prediction in these programs to make selections. Phenomic prediction provides an alternative opportunity that does not require the recurring costs of DNA sequencing. This research aimed to compare genomic and phenomic prediction in a diverse panel of soybeans. We found that phenomic prediction had the highest accuracy for seed yield, while genomic prediction was the best model for seed protein and oil. Combining genomic and phenomic data did not improve the predictive ability of the models.

Core ideas

- In-season phenomic prediction is able to outperform genomic prediction for seed yield.
- Genomic prediction outperformed in-season phenomic prediction models for seed protein and oil.
- Overlapping spectral indices were identified as the most predictive for seed yield, protein, and oil.
- Fusion of genomic and phenomic prediction did not increase the predictive ability of the combined models.

## 1 INTRODUCTION

Soybean [*Glycine max* (L.) Merr.] is one of the most important oilseed crops in the world and is of economic value because of its major source of both protein and oil. Soybean serves as an important resource in renewable energy production (Durrett et al., 2008), as well as in human and livestock nutrition (Singh et al., 2008; Ibáñez et al., 2020). Soybean breeding programs have worked to improve the crop’s genetic potential, with a historical focus on seed yield. Seed composition traits such as protein has been less of a focus in breeding for commodity-type soybeans, and the simultaneous improvement of both seed yield and protein is difficult due to the negative correlation between seed protein concentration and seed yield (Assefa et al., 2018; Singh et al., 2021b). The rising interest in renewable diesel is also causing a renewed breeding interest in seed oil (Egli, 2023). Seed yield, protein, and oil are quantitative traits controlled by many small-effect quantitative trait loci (QTL). Genome-wide association studies (GWAS) allow for a greater understanding of the genetic basis of these complex traits, and it has been successfully applied in soybean for seed yield (Ravelombola et al., 2021; Ayalew et al., 2022), and seed composition (Hwang et al., 2014; Zhang et al., 2018, 2019, 2021; Jin et al., 2023). However, the QTL often found in GWAS have minor effects, which makes them inefficient for use in marker-assisted selection (MAS) (Bernardo, 2008). An approach to assist in the selection of these complex traits is genomic selection (GS) (Meuwissen et al., 2001).

Genomic selection is a modern approach in plant breeding that utilizes genomic information to predict the performance of individual plants or lines (Crossa et al., 2017). Genomic selection involves genotyping and phenotyping a training population for estimating marker effects, which are then used to predict trait performance in breeding lines that are genotyped but not phenotyped. These predictions can then be used to make selections. GS has gained attention and adoption in soybean breeding due to its potential to accelerate the breeding process by reducing the time to complete breeding cycles and enhancing genetic gain (Zhang et al., 2016). In soybean, numerous studies have shown the usefulness of GS for predicting and selecting agronomic traits. An early soybean GS study demonstrated the application of GS for seed yield, plant height, and maturity (Jarquín et al., 2014). As a follow up study, GS was compared with traditional visual and end-season seed yield selections in untested populations and it was found that GS was as effective in increasing seed yield gains as the end-season seed yield selections (Bandillo et al., 2023). In another study, GS was found to be more accurate than MAS in predicting seed weight (Zhang et al., 2016). Previous reports have also demonstrated the usefulness of GS for seed yield with seed protein and oil traits within an active soybean breeding program (Stewart-Brown et al., 2019; Miller et al., 2023). The prediction accuracy of GS models heavily relies on accurate phenotypic information. Additionally, while genomic-assisted breeding methods are powerful and useful, the price point for genotyping can be a significant limitation for small-scale breeding programs (Wartha and Lorenz, 2021). An alternative or supplementary method to GS is the application of phenomic-assisted breeding methods, which are more amenable to different price points (Furbank and Tester, 2011). Genomic selection pipelines and phenomics-assisted methods both require accurate phenotyping to train and develop accurate models. However, acquiring large amount of accurate phenomics data has been a bottleneck in plant breeding as it is often laborious, subjective to human error, and destructive to plants (Furbank and Tester, 2011; Yang et al., 2020). This has resulted in the advancement of phenomics, with more accurate and rapid phenotyping of plants.

High-throughput phenotyping (HTP) is a set of techniques and technologies designed to measure plant traits rapidly and accurately on a large scale and has been extensively applied in soybean (Singh et al., 2021a). HTP aims to automate and streamline the phenotyping process, thereby reducing the labor and time required for evaluating plant characteristics (Gill et al., 2022). The ability of HTP to get precise traits and reduction in labor time increased the effectiveness of selection accuracy and, in turn, increased the genetic gain (Reynolds et al., 2020). HTP has been successfully implemented to measure and select numerous traits in soybean, such as tolerance to abiotic (Naik et al., 2017; Peirone et al., 2018; Dobbels and Lorenz, 2019; Zhou et al., 2020, 2021; Jones et al., 2024) and biotic stress (Nagasubramanian et al., 2019; Hatton et al., 2020; Rairdin et al., 2022). Seed yield has also been indirectly selected via HTP methods, such as in using crop canopy area (Moreira et al., 2020) and spectral reflectance (Parmley et al., 2019; Zhu et al., 2021).

HTP generates vast amounts of data encompassing various modalities such as images, sensor data, genomic information, and environmental parameters (Singh et al., 2016, 2021a). These multimodal datasets provide vast amounts of information about plant traits; however, analyzing such complex data is difficult, and traditional statistical methods are not always able to capture the complex relationships between phenomic predictors and traits of interest (Danilevicz et al., 2022). Machine learning (ML) techniques are viable tools for such datasets (Singh et al., 2018). ML algorithms can automate the analysis of phenotypic traits from various datasets. ML models can be used to predict plant traits, or phenomic prediction (PP), based on multimodal data inputs, including genomic information, environmental factors, and historical phenotypic data (Shook et al., 2021a). Models such as partial least squares regression (PLSR) and random forest (RF) have been found to have moderate predictive ability for seed yield based on canopy spectral reflectance data (Christenson et al., 2016; Parmley et al., 2019; Chiozza et al., 2021). Extreme gradient boosting (XGBoost) is another method that has been found to have moderate predictive ability for soybean seed yield, protein, and oil on a field and county-level prediction scale (Torsoni et al., 2023; Hernandez et al., 2023), although its predictive power on breeder plots is not well studied (Carroll et al., 2024).

Early, non-destructive, and real-time prediction of complex traits such as seed yield, protein, and oil on a plot scale has several potential uses in a breeding program, such as for early-season selection and harvesting decisions in breeding programs (Moreira et al., 2019; Singh et al., 2021b), and is a useful feature for developing a cyber-agricultural system which provides a framework to integrate computation and decision making in a breeding program (Sarkar et al., 2024). Both genomic prediction (GP) and phenomic prediction (PP) have the potential to be used for early season predictions, enabling earlier selections in breeding programs.

In our study, we compared the usefulness of genomic and phenomic prediction using a diverse soybean panel. We conducted GWAS to detect QTL for these traits and compared the predictive ability to select via genomic or phenomic tools. We report on the results from the GWAS that compared multiple methods applied to a soybean diversity panel. We then report on the prediction accuracy of individual GP and PP models that were trained on the soybean diversity panel. We compare the prediction accuracy and usefulness of these models in three major soybean traits: seed yield, protein, and oil. Finally, we assessed the improvement in the prediction accuracy when we added phenomic covariates and SNPs to the different GP and PP models, respectively.

## 2 MATERIALS AND METHODS

### 2.1 Germplasm

A panel of 292 diverse soybean accessions from 19 different countries of origin was utilized. These accessions were in the maturity groups of MG I to MG III and were previously selected in the field for their absence of lodging and shattering. All accessions are from either the USDA Soybean Germplasm Collection or are parents of the Soybean Nested Association Mapping (SoyNAM) panel, for a total of 260 and 32 accessions, respectively. The 260 accessions from the USDA collection are plant introductions (PIs) that have publicly available genotype data from the SoySNP50K BeadChip (Song et al., 2015). The 32 accessions from the SoyNAM parents are a mix of elite and diverse lines, with the elite being breeding lines released from public breeding programs and diverse lines being crosses of elites and PIs.

### 2.2 Experimental Design

As outlined by Parmley et al. (2019), the experiment was conducted in 2016 and 2017. In 2016, two locations were grown in Boone County, Iowa (AGRON16), and Cass County, Iowa (LEWIS16). Four locations were grown in 2017, with the two previous locations from 2016 (AGRON17 and LEWIS17) and two new locations in Boone County (MARS17) and in Monona County, Iowa (CAS17). This resulted in six unique environments (location-year combinations). Each environment was planted in a unique alpha lattice design, which consisted of two replications and 30 incomplete blocks. Plots consisted of four rows at a length of 4.6 m and a seeding density of 296 K seeds ha^-1^.

### 2.3 Phenotypic Measurements

#### 2.3.1 Field Based

Plots in each environment were phenotyped for phenomic traits at two time points. These time points were made when the plots were at the same approximate growth stages and were S1: flowering (R1-R2) and S2: pod set (R3-R4) (Fehr et al., 1971).

Canopy spectral reflectance was measured using a FieldSpec 4 Hi-Res (ASD Inc., Boulder, CO, USA) spectroradiometer. The spectroradiometer covers 2150 spectral wavebands within the range from 350 to 2500 nm in one nm bands. Data was collected by positioning the fiber-optic cable 1.0 m above the canopy in a nadir position as recommended by the manufacturer. In 2016, height was maintained via a tripod, and in 2017, a phenotyping cart was utilized to maintain the height. Two reflectance measurements from each of the middle two rows of the plot were collected. Measurements were taken on cloudless days within two hours of solar noon. The spectroradiometer was calibrated every 20 minutes during data collection by normalizing it to a white reference panel (Specralon, Labsphere Inc., North Dutton, NH, USA).

The waveband data was processed as previously outlined in Parmley et al. (2019). Briefly, the mean reflectance of the plot was calculated by averaging the two observations. The repeatability of each waveband was calculated across locations, and wavebands with a repeatability below 0.3 were removed. From the remaining wavebands, the mean reflectance across 10 nm blocks was calculated to produce 178 average wavebands. Vegetation indices (VI) were calculated from the individual wavebands to get VIs that have been previously associated with physiological traits (Table S1).

Seed yield (SY, kg ha^-1^) was measured by harvesting the middle two rows of the plot using an ALMACO SP20 combine once all plots had reached physiological maturity (R8). Seed moisture was measured on harvested plots and used to adjust plot SY values to 13% moisture.

#### 2.3.2 Post Harvest

In 2021, 100 grams of remnant seed samples were sent to the University of Minnesota. Seed from two environments, AGRON16 and MARS17, were not sent for NIR due to insufficient remnant seed sources. Samples were ground to 1mm in particle size and subsequently scanned with a DA 7250 (PerkinElmer Inc., Wellesley, MA, USA) near-infrared reflectance (NIR) machine to predict seed protein and oil on a dry matter basis using a global calibration set provided by PerkinElmer Inc.

### 2.4 Statistical Analysis

Best linear unbiased predictions (BLUPs) and estimates (BLUEs) were calculated for each line using the H2cal function from the inti package (Lozano-Isla, 2023) in R. A mixed linear model was fit with the following equation:

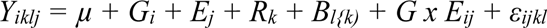

Where *Y* is a vector of observed phenotypes, *µ* is the grand mean, *G_i_* is the effect of the *i*th genotype, *E_j_* is the effect of the *j*th environment, *R_k_* is the effect of the *k*th replicate, *B_l{k)_* is the effect of the *l*th incomplete block nested in the *k*th replicate, *G x E_ij_* is the interaction effect of genotype and environment, and *ε_ijkl_* is the residual error.

Analysis of variance (ANOVA) for seed yield, protein, and oil was conducted to evaluate the effect of genotype, environment, and their interaction as fixed terms and the remaining terms as random using a mixed linear model in the lme4 package in R. This model included the same term as the model used to calculate BLUPs. ANOVA assumptions were tested using the Shapiro-Wilk test for normality and Bartlett’s test. Outliers were removed by calculating the studentized residuals for each observation of each trait and excluded from the analysis with values ± 3.78 (Lund, 1975).

Broad sense heritability (*H^2^_Piepho_*) was calculated via the H2cal function in the inti package and was determined by the equation:

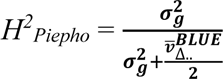

Where 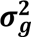 is the genotypic variance and 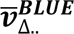 is the average standard error of the genotypic BLUEs.

Additionally, the SNP-based heritability (*h^2^_SNP_*) (Yang et al., 2017) was calculated. Genotypic data for 268 of the 292 genotypes was obtained from the publicly available SoySNP50K BeadChip dataset (Song et al., 2015). SNP markers with a missing rate of >10% were removed from further analyses, and the remaining missing SNPs were imputed using the Expectation-Maximization (EM) method in the A.mat function from the rrBLUP package (Endelman, 2011) in R. Unlike conventional estimates of heritability, the A matrix is used to calculate marker-based genetic variance (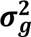) associated with the genotypes. The *h^2^_SNP_* is computed using the following equation:

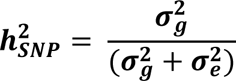

Where 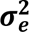 is the residual variance. The R package heritability (Kruijer et al., 2023) was used to compute the SNP-based heritability using the marker_h2 function.

### 2.5 Genome-Wide Association Study & Genomic Prediction

GWAS was performed in three different programs: TASSEL (Bradbury et al., 2007), GAPIT v3 (Wang and Zhang, 2021), and SVEN (Li et al., 2023b). The same phenotypic dataset of BLUEs was supplied to each of the GWAS methods. The previously mentioned subset of 268 genotypes was used due to available Soy50KSNP data. Accessions not in the public dataset were dropped from all further GWAS and GP analyses. Of the panel of 292 genotypes, 268 accessions had Soy50KSNP data publicly available for use (Song et al., 2015). The SoySNP50K dataset has 47K markers across the genome, which was imported into TASSEL and filtered on a minor allele frequency (MAF) of 1%, leaving 38K markers for further GWAS analysis. Genotypic data was imputed and used to calculate the kinship matrix with a centered identity by state (IBS). A principal component analysis (PCA) with four components was conducted to account for any familial or population structure. The numerical genotype, kinship matrix, and PCA were exported from TASSEL for further use in GAPIT and SVEN. The FarmCPU (Kusmec and Schnable, 2018) and BLINK (Huang et al., 2019) models were selected for use in GAPIT. For both the selected models and TASSEL, a MLM model was used, which is described as:

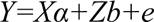

Where *y* is a vector of phenotypic observations, *α* is a vector of fixed effects that include the population structure, *b* is a vector of the genetic effects defined by a kinship matrix, *X* and *Z* are design matrices, and *e* is the vector of residual effects. A LOD threshold of 5.87 was selected (FDR < 0.01) for detecting significant marker trait associations (MTAs) across TASSEL and the selected GAPIT models. The Selection of Variables with Embedded Screening (SVEN) is a Bayesian method based on a hierarchical multi-locus model. SVEN was utilized using a prior lambda shrinkage of 20, and a marginal inclusion probability (MIP) of 0.5 was used to select significant MTAs.

A genomic best linear unbiased prediction (GBLUP) model was implemented using the rrBLUP package (Endelman, 2011). Genotype BLUPs data from individual environments and from across environments were randomly divided into training (80%) and validation (20%) sets. A ten-fold cross-validation (CV) was performed on the training set to avoid inflated estimates of predictive ability. The cross-validation was repeated 100 times, with each repetition randomly re-sampling the training set. The phenotypes of the validation set were masked in the model training. A prediction accuracy denoted as genomic prediction accuracy (GPA) was calculated as the correlation between the observed genotyped BLUP based on phenotypic data and the prediction of the genotypic effect from the GBLUP model. Both Spearman rank correlations and Pearson correlations were calculated.

### 2.6 Phenomic Prediction

Three different machine learning model architectures were trained for use in predicting seed yield, protein, and oil using VIs. Model architectures were separately trained using all the available VIs. The caret package (Kuhn, 2008) in R was implemented for model training and hyperparameter tuning of all models. Three different methods were used for model training: partial least squares regression (PLSR), random forest (RF), and extreme gradient boosting (XGBoost), and were compared in their prediction accuracy. A repeated cross-fold validation with five repeats and ten folds was used to gauge model performance during training. The variable importance of each model was calculated using the varImp function.

For cross-validation, a split of 80% of accessions was included in model training, while the remaining 20% was kept unseen for use in the validation set. This split was used for both the across-environment scenario and for the individual environment scenarios. Model prediction performance was calculated using the Spearman rank correlation coefficient between the observed values and the predicted values of the validation set. This rank correlation is practical in a breeding program that seeks to rank the performance of untested genotypes. Each model was run ten times and reported correlations and feature importances were averaged from the ten runs.

## 3 RESULTS AND DISCUSSION

### 3.1 Phenotypic Performance

Seed yield was measured on 3504 plots across six environments. The mean seed yield observed across the 292 accessions was 2126 kg ha^-1^. The PI germplasm had the lowest seed yield (1974 kg ha^-1^), followed by the diverse (3624 kg ha^-1^), and the elite germplasm had the highest seed yield (4084 kg ha^-1^). The broad sense heritability estimates were high, indicating low variation due to the environment or residual sources, as well as a high amount of genetic variation in the measured panel. However, the SNP-based heritability was lower and varied across environments, with the lowest value occurring for the combined environment analysis.

Seed protein and oil were measured on 1866 plots across four environments and are presented on a dry weight basis. The mean seed protein was 41.82%, with the PI group having the highest mean (41.93%), followed by the diverse (39.76%) and the elite group having the lowest mean seed protein content (39.36%). Mean seed oil was 18.6%, with the PI group having the lowest mean seed oil content (18.41%), followed by the diverse (20.76%), and the elite group having the highest mean seed oil content (20.9%). Seed protein and oil both had moderately high estimates for broad sense and SNP-based heritability, with oil having higher estimates. As seen with yield, the SNP-based heritability estimate was often lower in the combined environment analysis, which could be due to the variation between the environments. For all traits, the PI group had the most extensive variation, while the elite and diverse groups largely overlapped (Figure 1). A further breakdown of the descriptive statistics for each trait can be found in Table 1.

**Figure 1.**
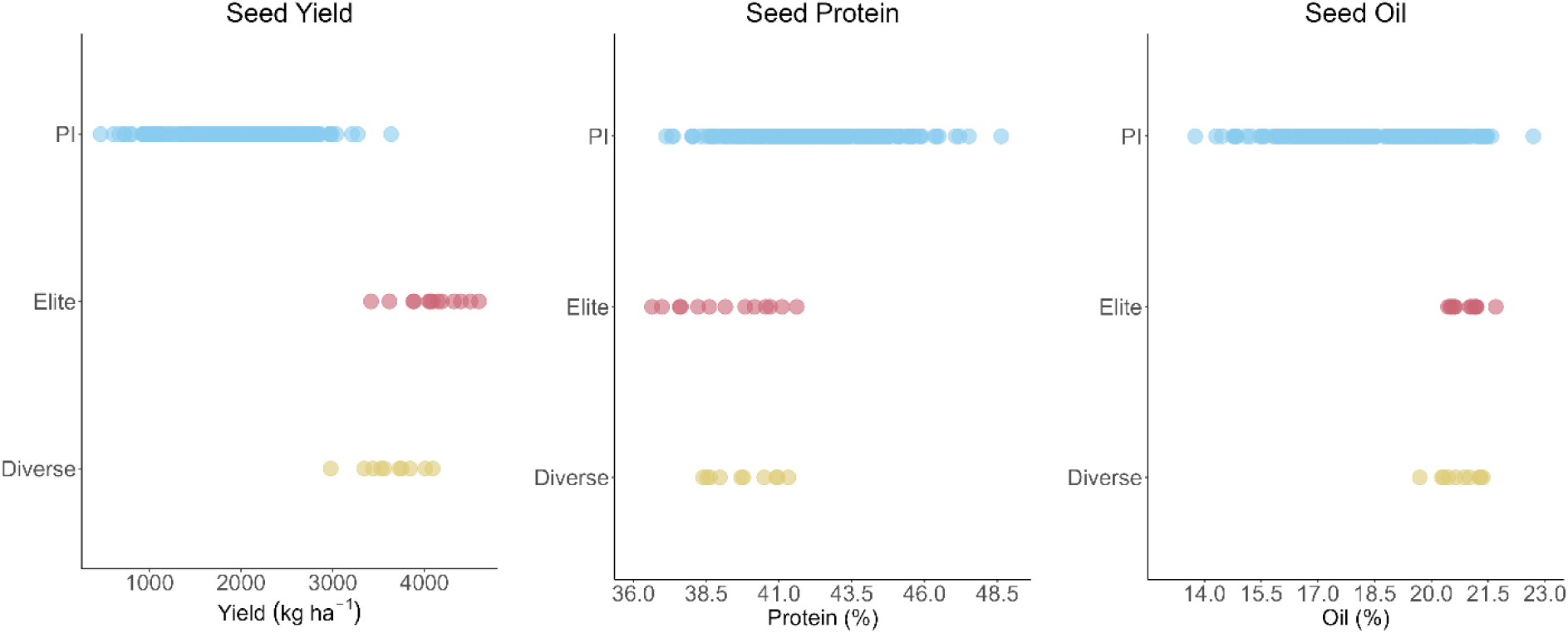
Seed yield, protein, and oil of 292 genotypes grouped as PI (n=269), elite (n=13), and diverse (n=10)

**Table 1.**
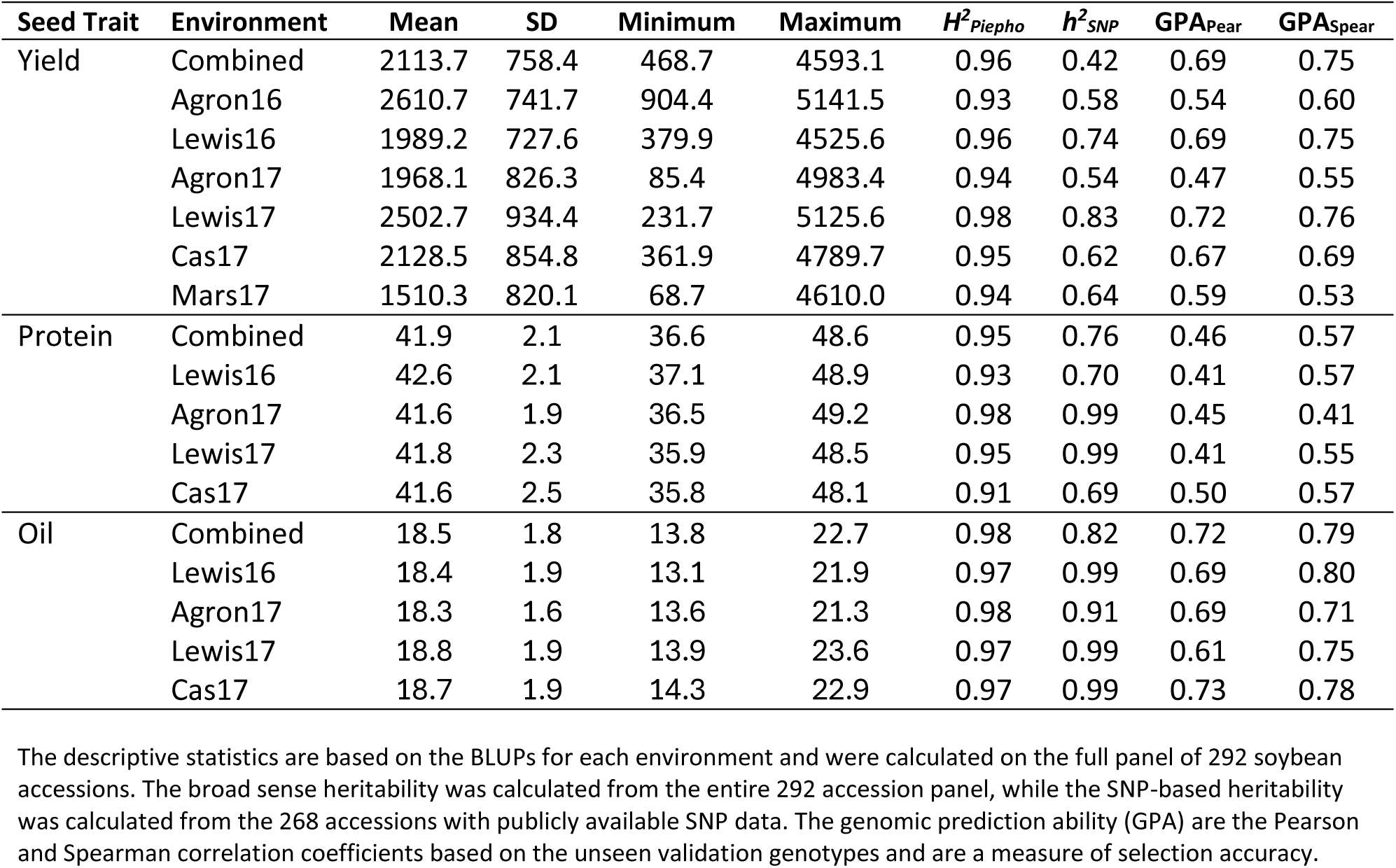
Summary of the descriptive statistics, broad sense heritability, and SNP-based heritability for yield, protein, and oil.

### 3.2 Genome-Wide Association Study

Across the four GWAS methods, there were 186 MTAs detected as significant across the different traits and environments. Due to overlapping MTAs across GWAS methods, traits, and environments, there was a total of 127 unique MTAs identified (Table S2). Figure 2 demonstrates that many MTAs were identified across the four GWAS methods, and that all programs have some agreement in reporting. The distribution of significant MTAs (Table S3) identified multiple MTAs in a single trait that were shared across GWAS methods and across different environment analyses. FarmCPU detected the greatest number of significant MTAs, followed by SVEN, BLINK, and TASSEL. Seed yield was the trait with the highest number of unique SNPs detected, with a total of 56 unique, significant MTAs detected across the environments and GWAS methods. Seed oil had the second-highest number of unique significant MTAs (= 51). Seed protein had the lowest number of significant MTAs detected (= 21). No MTAs detected for seed yield overlapped with protein or oil MTAs. One SNP, *ss71562177*, was detected for both seed protein and oil, suggesting a pleiotropic effect. Multiple MTAs for each trait were detected in more than one environment, suggesting a SNP with greater stability across different environments. In two examples with seed yield associated MTAs, neighboring MTAs on chromosomes 7 and 15 were associated with seed yield in different environments, indicating regions of interest.

**Figure 2.**
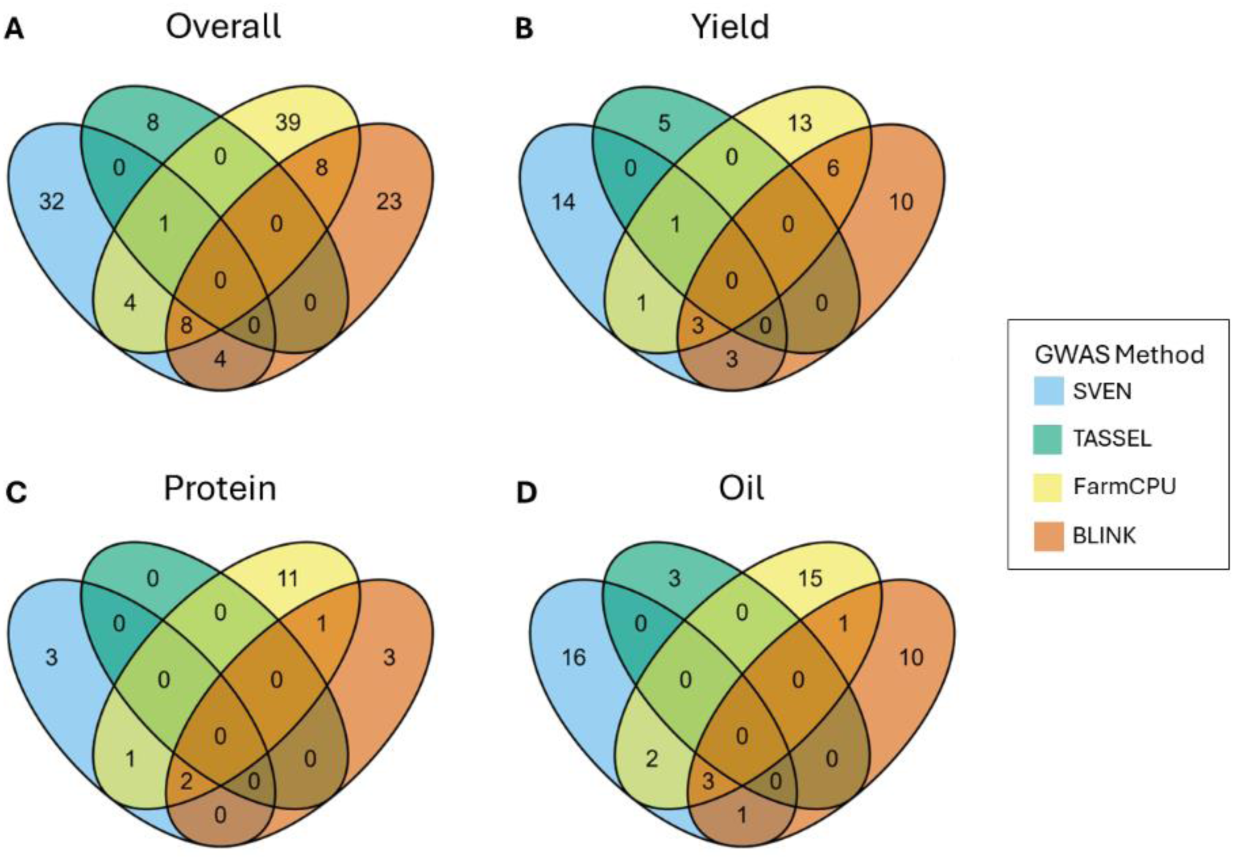
Venn diagrams showing the number of significant SNP markers grouped by the GWAS method. (A) shows the overlap of markers across all traits, (B) is the overlap for yield associated SNPs, (C) is the overlap for protein associated SNPs, and (D) is the overlap for oil associated SNPs.

#### 3.2.1 Seed Yield

A total of 56 associated MTAs were detected across the GWAS methods and locations for seed yield. Eleven MTAs were found in the combined environment analysis (Table S2), with four of these MTAs also detected in at least one of the individual environment analyses, indicating the stability of these MTAs across environments. Two regions of interest were identified for seed yield in this study on chromosomes 15 and 16. The region on chromosome 15 had three different SNPs detected (*ss715620226*, *ss715620227*, and *ss715620229*) in a 32.2 kbp region. The three MTAs were found to be associated in three individual environments as well as in the combined environment analysis and were detected by three of the different GWAS methods. Within the region of these three MTAs are three candidate genes, *Glyma.15G130800*, *Glyma.15G130900*, and *Glyma.15G131200*. These genes are annotated as a GTP binding protein, a leucine-rich receptor-like kinase protein, and a 6-phosphogluconate dehydrogenase family protein, respectively.

#### 3.2.2 Seed Composition

A total of 21 unique MTAs were associated with seed protein content across the GWAS methods and environments. Of these MTAs, 12 were associated with the combined environment analysis. Three SNPs, *ss715615568*, *ss715627800*, and *ss715635419*, were detected in the combined environment analysis as well as in multiple individual environments. The SNP *ss715615568* on chromosome 13 was found in three individual environments and the combined, with all methods except TASSEL detecting this MTA in the combined analysis. The second SNP, *ss715627800*, was found in two of the individual environments along with the combined environments. The third SNP, *ss715635419*, is located on chromosome 19 and was associated in two individual environments and the combined environment.

For seed oil content, 51 unique MTAs were associated across the methods and environments, with 17 of these MTAs being associated with the combined environment analysis. The SNP *ss715591697* is located on chromosome 5 and was detected in one individual environment and the combined environment analysis. Another MTA of particular interest is *ss715621777*, which is located on chromosome 5. This MTA was associated with seed protein in a single environment and was associated with seed oil in two individual environments and the combined environment analysis, indicating a stable pleiotropic effect.

### 3.3 Genomic Prediction

Genomic prediction accuracies (GPAs) for all three traits and environments are shown in Table 1. The means differed between the traits, with seed oil having the highest GPA across environments while seed protein had the lowest GPA. In most of the individual environment predictions, the GPA was lower than the GPA for the predictions across environments. This trend was true across traits and across the testing and validation sets.

### 3.4 Phenomic Prediction

Three separate ML models were trained for predicting seed yield, protein, and oil based on VIs. Seed yield had the highest predictive ability based on rank correlations (0.83; p < 0.01), while seed protein had the lowest correlations (0.50; p < 0.01). Given the early reproductive stages that measurements were taken (flowering and pod set), it is expected that the predictive ability for seed composition traits would be lower. For all three predicted traits, random forest had the highest predictive ability, followed by XGBoost for the seed yield and oil traits. The phenomic prediction had a higher rank correlation for seed yield than the GBLUP genomic prediction model (Figure 3), although the phenomic prediction model did not outperform GBLUP for the seed protein and oil traits.

**Figure 3.**
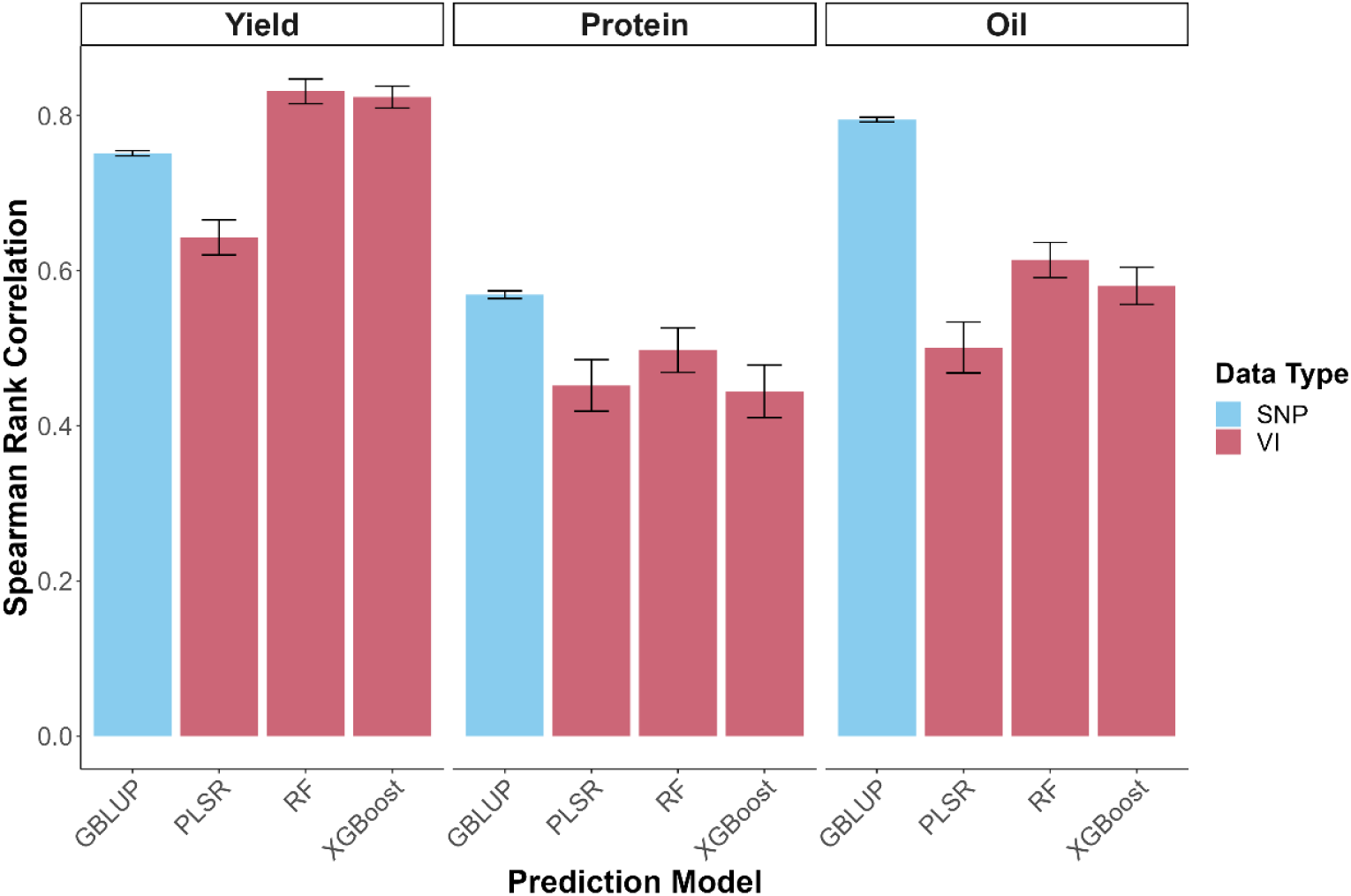
Spearman rank correlations were obtained from different prediction models (1) GBLUP genomic prediction, (2) partial least squares regression (PLSR), (3) random forest (RF), and (4) eXtreme gradient boosting (XGBoost). The genomic prediction model was trained on the Soy50KSNP chip with 268 genotypes. The three phenomic prediction models were trained on vegetation indices for predicting yield, protein, and oil on 292 soybean genotypes. Error bars represent the standard error around the mean.

The feature importance analysis indicated that the type of prediction model changed the importance of the features. Feature importance also varied across the predicted trait. From the 10 VIs at two time points, 14 of the combinations appeared in the top 5 of a model-trait combinations for the across-environment analysis. Eight of these VIs came from the S2 timepoint, while the remaining six were from the S1 timepoint. The most common VI in the top five for importance across all traits is VREI2. Other commonly recurring VIs include PRI, RARSb, and RARSc. A summary of the top five VIs for each trait and model can be found in Figure 4.

**Figure 4.**
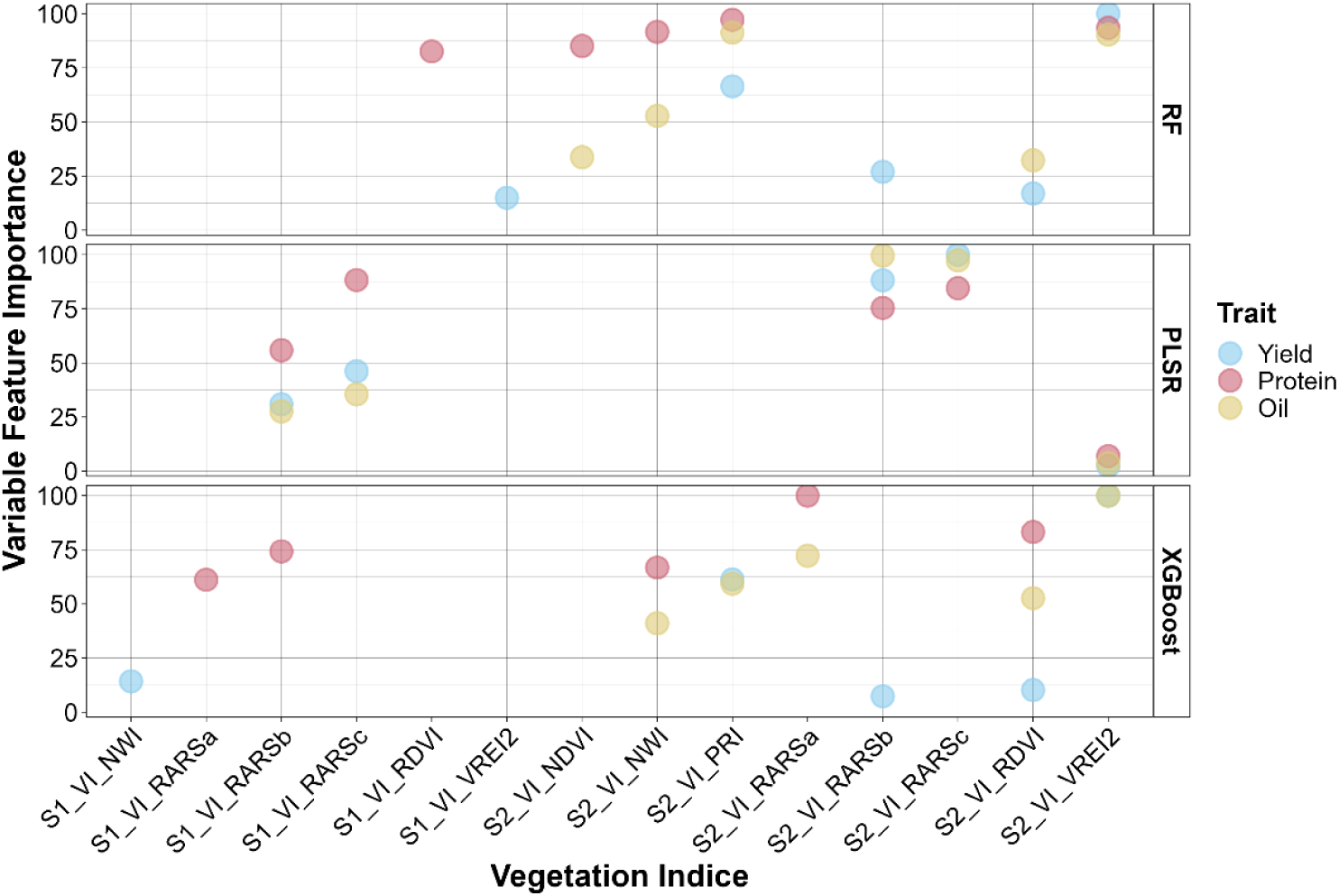
The top five timepoint and VI combinations featured in the average variable importance for each trait and model in the combined environment analysis.

### 3.5 Combining Genomic and Phenomic Data for Predictions

We selected the top five features from the RF models of each trait to be included as covariates in the GBLUP model. These features were selected as the RF models had the greatest Spearman rank correlation (Figure 3). Additionally, the SNPs detected as significant for the across-environment GWAS were used as additional features for prediction in a new RF model. As seen in Figure 5, the addition of selected VIs in the GBLUP model did not change the rank correlation as compared to the genomic-only GBLUP model. Similarly, the addition of selected SNPs to the RF model did not significantly change the correlations from the original VI-only model.

**Figure 5.**
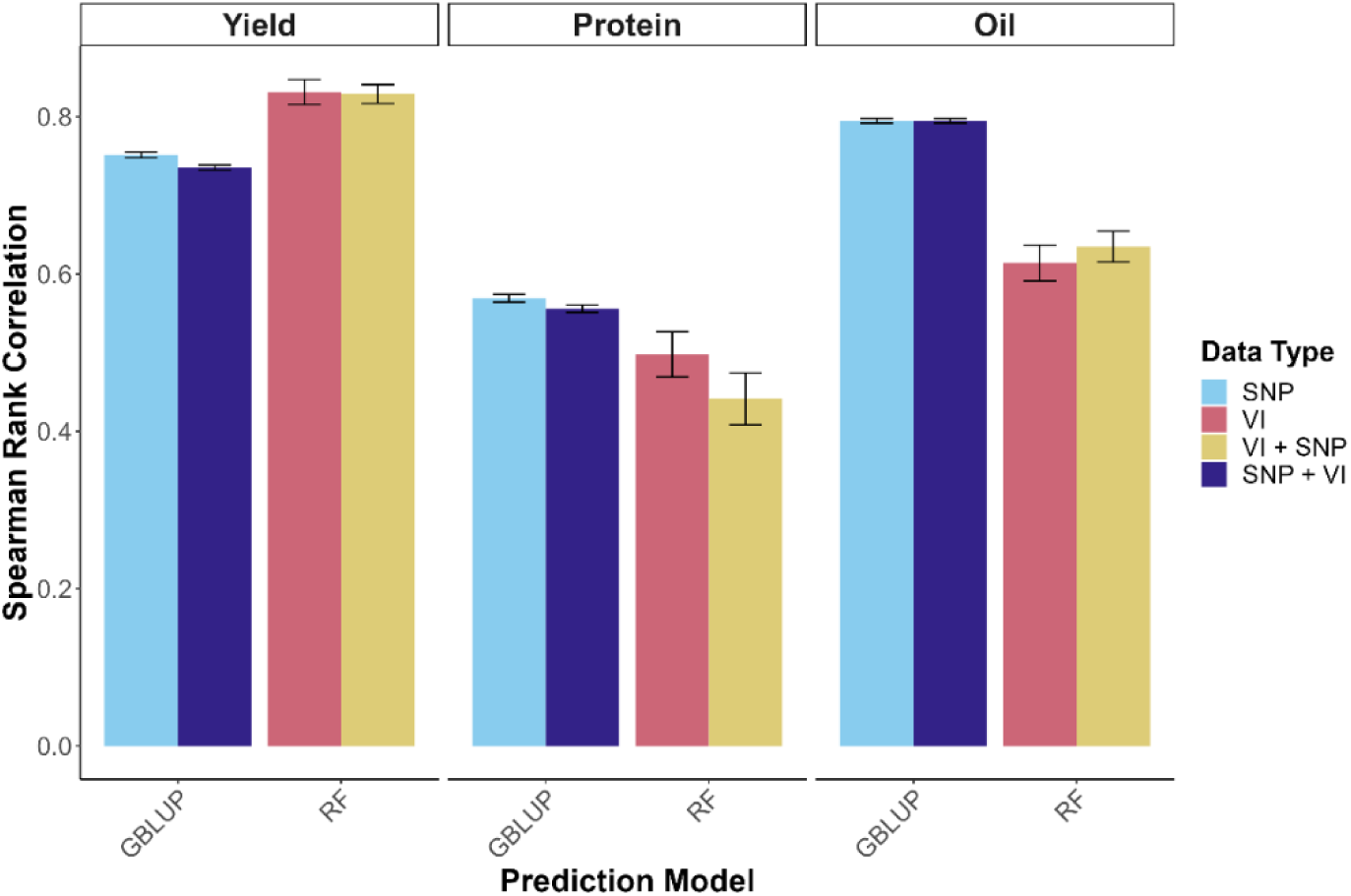
Spearman rank correlations were obtained from GBLUP (genomic prediction) and from the random forest (phenomic prediction). The GBLUP model was trained on genomic data only (SNP) and with the top five VI as covariates (SNP + VI). The RF model was trained with all VIs (VI) and with a subset of SNPs selected from GWAS results (VI + SNP). Error bars represent the standard error around the mean.

## 4 DISCUSSION

Seed yield, protein, and oil are primary focus areas for soybean breeders. Soybean breeders and geneticists aim to utilize diverse genetic accessions in the development of improved soybean cultivars. Tools such as marker-assisted selection, genomic selection, and phenomic selection are useful and have the potential to increase the use of diverse genetic material in soybean cultivar development.

### 4.1 Phenomic Performance and Heritability Estimates

Unadapted soybean genotypes contain an extensive range of seed protein and oil contents, as well as variation for seed yield. This variation often contains genes of interest that could be used to improve these traits for the development of new soybean cultivars (Carter et al., 2004). We noted PIs that would be useful for specialty breeding of high protein or high oil lines (Table S4). The H^2^ estimates of these traits were high, which could indicate a low environmental variance due to similar environments. The h^2^_SNP_ estimates, which are closer to estimates of narrow-sense heritability, were lower than the H^2^ estimates, which is expected due to the loss of the dominance and epistatic variance contributions. This indicated that while these traits are complex with multiple gene interactions and interaction with the environment.

### 4.2 GWAS and Candidate Genes for Soybean Seed Yield and Composition

In this study, we report on GWAS QTL from a diversity panel of soybean accessions. Four different GWAS methods were used, with the programs FarmCPU, BLINK, and SVEN having the greatest amount of overlap between methods, which is similar to previously reported studies (Rairdin et al., 2022; Chiteri et al., 2023). A total of 56, 21, and 51 MTAs were significantly associated with seed yield, protein, and oil, respectively, across the environments and GWAS methods. A total of 11, 12, and 17 MTAs were found to be associated in the combined environment analysis for seed yield, protein, and oil.

In the present study, several of the associated MTAs were previously reported loci. Two of the MTAs associated with seed yield in the combined environment analysis, *ss715624192* and *ss715624199*, have been previously detected for seed yield and shattering in a single environment in Kansas (Ayalew et al., 2022). The SNP *ss715630114*, which was associated with seed yield in the combined environments, is 98 kbp away from an MTA previously reported for seed yield across three years in China (Ravelombola et al., 2021). Two additional seed yield associated MTAs, *ss715624379* and *ss715626802*, are near previously reported MTAs. The first is 47.4 kbp away from a MTA associated with seed yield (Ayalew et al., 2022), and the second MTA is within a QTL detected for biomass in drought conditions (Li et al., 2023a).

The MTAs associated with seed protein or oil content also had several previously reported regions. The most prominent of the previously reported regions is the SNP *ss715621777*. This MTA was associated with seed oil in several individual environments and the combined environment analysis, as well as seed protein in a single environment. This MTA has been previously reported for seed protein and oil (Bandillo et al., 2015), seed composition development (Zhang et al., 2018), and is within a QTL previously detected for seed protein content and seed oil content (Qin et al., 2022; Jin et al., 2023; Clevinger et al., 2023). A deletion was identified in the coding sequence of the nearby candidate gene *Glyma.15G049200*, which is a sugar transporter *GmSWEET39* gene, indicating that this is the causative gene for the numerous reported effects on seed composition (Zhang et al., 2020).

### 4.3 Genomic Prediction in Soybean

We assessed the prediction accuracy for seed yield, protein, and oil content for use in scanning diverse soybean accessions. The quantitative nature of these traits makes genomic prediction a promising tool for use in soybean breeding programs. Previous studies have reported on the GPA of genomic selection for use in soybean seed yield, protein, and oil. We report a GPA across environments of 0.75, 0.57, and 0.79 for seed yield, protein, and oil, respectively. This estimate of accuracy for seed yield is towards the higher end of the previously reported range of 0.26-0.74 (Jarquín et al., 2014; Duhnen et al., 2017; Matei et al., 2018; Đorđević et al., 2019; Stewart-Brown et al., 2019; Beche et al., 2021; Ravelombola et al., 2021). Studies on the application of genomic prediction for soybean seed protein and oil are less prevalent than for seed yield. Our estimated GPA for seed protein is within the range of previously reported GPA across populations, which ranges from 0.45-0.81 (Duhnen et al., 2017; Stewart-Brown et al., 2019; Beche et al., 2021; Qin et al., 2022; Sun et al., 2022). The predictive ability for seed oil is previously reported on less frequently than reports on seed protein, although our estimate is in line with those previous reports, which have GPAs ranging from 0.71-0.85 (Stewart-Brown et al., 2019; Beche et al., 2021; Sun et al., 2022). The GPA for the individual environments was generally lower for all three traits, which was also noted in the broad sense heritability estimates. This could be due to a greater environmental effect being present in a single environment. We acknowledge the reported GPA is likely inflated when compared to GS models deployed in elite breeding germplasm. However, these models could help in front-loading useful genetic diversity for breeding programs.

### 4.4 Phenomic-Enabled Prediction

During the early and mid-stages of testing, breeders are often more interested in identifying the top-ranking experimental lines rather than predicting the actual seed yield and performance. We used the Spearman-rank correlation coefficient to assess the model performance and to identify models with high-ranking accuracy to be used as a selection tool for breeders. We observed that VIs produced modest prediction accuracies for seed yield which were higher than the prediction accuracies for the GP model.

The model with VIs was moderately accurate in predicting seed protein and oil. In comparison to the prediction accuracy with the GBLUP model, the VI prediction model had a higher rank correlation coefficient for seed yield, which is similar to results found in maize (Adak et al., 2023; DeSalvio et al., 2024) and wheat (Dallinger et al., 2023). Postharvest NIR of seeds has also been found to result in phenomic prediction accuracies similar to or greater than genomic prediction in wheat (Robert et al., 2022) and soybean (Zhu et al., 2021), although the utility of this is limited given the measurements must occur post-harvest. The higher accuracy for seed yield via the PP model could be due to the time series VIs capturing in-season stress responses, which affected the seed yield. The genomic prediction method is more static and cannot respond to the in-season changes to the environment in comparison. The genomic prediction model outperformed the phenomic VI model for seed protein and oil. This could be because canopy reflectance was measured in the early reproductive stages. During this time, seeds are starting to develop, the seed number is not yet set, and assimilates have not yet been partitioned into the seeds (Rotundo et al., 2009). Additionally, soybean pods develop under the cover of the canopy, making it hard to measure the seeds during development directly. While seed yield is also developing under the canopy, crop yield determination has been found to start earlier based on the amount of intercepted photosynthetically active radiation (iPAR) (Monteith, 1972). This has resulted in models using hyperspectral wavelengths to better predict soybean seed yield compared to seed protein and oil (Chiozza et al., 2021). In future studies, the combination of spectral data and genomic prediction in complex models would be interesting as they are complementary to each other, as previously shown in wheat (Crain et al., 2018; Sandhu et al., 2021). Genomic prediction has also been found to have improved prediction accuracy in soybean seed yield by including canopy coverage data (Howard and Jarquin, 2019).

Among the VIs, VREI2 was the most commonly featured VI in the top five feature importance. This VI had the highest feature importance for seed yield when using the RF and XGBoost models and was also the second highest-rated feature in these models for seed protein and oil. VREI2 was previously reported as useful in soybean seed yield phenomic prediction (Parmley et al., 2019), and is associated with canopy area, chlorophyll concentration, and leaf water content (Vogelmann et al., 1993). Other notable VIs include PRI, RDVI, RARSb, and RARSc. These VIs all appeared five or more times as top features in the variable importance analysis, regardless of time point. These VIs are related to the photosynthetic radiation use efficiency (Gamon et al., 1997), vegetation coverage (Roujean and Breon, 1995), chlorophyll b content (Chappelle et al., 1992), and carotenoid contents (Chappelle et al., 1992), respectively.

### 4.5 Combining Genomic and Phenomic Predictions

We were interested in utilizing both genomic and phenomic data to leverage the additional information for increased predictive ability. As shown in Figure 4, the addition of selected VIs as covariates in the GBLUP or of selected SNPs as additional features in the RF model did not increase the correlation coefficient. In a previous study in wheat, the addition of HTP data as covariates did not increase the correlation resulting from a GBLUP model, although a multi-variate GBLUP model with secondary traits did increase the predictive ability (Crain et al., 2018). This is in contrast to another study which found an increase in predictive ability for wheat grain yield and grain protein content by including VIs as either covariates or in a multi-variate GBLUP model (Sandhu et al., 2021). In soybean, information from canopy images has been found to increase the prediction accuracy of genomic prediction when using a multi-variate model (Howard and Jarquin, 2019). The lack of increase in the predictive ability of our models could be due to the high correlations we had with the simpler models, and a lower environmental effect in our dataset, as indicated by the high heritabilities.

As advances are made in HTP and ML, there is a greater opportunity to explore the application of different omics in soybean breeding, thus working to increase genetic gain (Singh et al., 2021a). As sequencing costs continue to decline, the application of GS becomes more feasible (Bhat et al., 2016), especially in public breeding programs. Additionally, the digitization of genomic data and historical breeding datasets in public databases is allowing for the training of larger and more robust GP models (Jarquin et al., 2016). Our results indicated the usefulness of a PP model for select traits in soybean breeding using VIs. However, this data was ground-based and can still be laborious to collect. The usefulness of predictors collected via drones for final soybean seed yield prediction using ML models has been demonstrated as well (Herrero-Huerta et al., 2020). The use of drones has also been found to be useful in phenomic prediction and supporting genomic prediction in crops including maize and wheat (Herr et al., 2023). Further prediction models should continue to investigate further the usefulness of remote sensing platforms for use in collecting accurate data to be used in both GP and PP models (Guo et al., 2021).

In addition to remote sensing, other sources of data should be investigated for any gains in prediction accuracy. Novel data sources should be investigated for their ability to capture phenotypic traits which may be useful covariates for prediction models. Such sources could include spoken phenotypic descriptions (Yanarella et al., 2024) and 3D point clouds to investigate the role of different plant organs (Young et al., 2023, 2024). Secondary traits could also be explored for addition to models based on their relationships with specific traits, such as nodulation with seed protein (Vollmann et al., 2011; Carley et al., 2023). Environmental and management data are two additional areas that could be further leveraged to train more robust prediction models. For example, soil properties have been previously reported to assist in capturing genotype-by-environment (GxE) interactions for soybean in both GP applications (Vieira et al., 2022) and for spatial adjustments in advanced line testing (Carroll et al., 2024). Weather data such as temperature and rainfall have also been found to be important predictor variables for soybean yields (Shook et al., 2021a). Integrating additional management data could provide greater predictive ability and allow for complex genotype-by-environment-by-management (GxExM) interactions to be captured in the prediction models for complex traits (McGowan et al., 2021). Additional data can be sources from multiple different sources, which can cause complexities in analysis and prediction models.

The further development of ML models can assist due to the ability to handle complex datasets and large amounts of data, as shown in applications in crop stress phenotyping (Singh et al., 2018) and for complex GP models (Grinberg et al., 2020). Advances in GP models are also being made continuously, such as including integration with crop growth models (Technow et al., 2015; Messina et al., 2018), tracing the source of parental alleles (Shook et al., 2021b), inclusion of GxE terms (Song et al., 2022), and multi-trait models (Dos Santos et al., 2020). Advances in GP models in animal breeding can also be leveraged and transferred to the plant breeding domain, with advances in Bayesian models (Wolc and Dekkers, 2022) and two-step models (Ni et al., 2017). All of these models can be used in advancing genetic gain, which has been found to need multiple models to fit the parameters of different breeding strategies (Krause et al., 2023).

## 5 CONCLUSION

Here, we showed the usefulness of genomic selection in soybean for three separate traits and found that predictions across multiple environments had the greatest predictive accuracy. Over the last decade, genomic selection has become instrumental in driving genetic gain for crop improvement, although the uptake in public breeding programs has been slower (Wartha and Lorenz, 2021). While genomic selection is a powerful tool, it is limited due to the high costs of the markers, the dependence of related individuals for training and applying the model, and the less dynamic nature for in-season predictions with time series data in a changing environment. With the advances in high throughput phenotyping, we demonstrated the use of VIs as a phenomic prediction tool. We found that for seed yield, the phenomic prediction method has the capacity to perform equal or better than the genomic selection method and does not require yearly genotyping of accessions. The phenomic prediction model was not able to perform as well as genomic selection for seed protein and oil. However, further studies with later season measurements to test the predictive ability of such seed traits are needed. Using the results from the GWAS, the addition of significant SNPs as predictors in the phenomic prediction model did not show any improvement in predictive ability. Phenomic selection adds another tool for use in cultivar development to increase genetic gain and could be used to complement genomic selection as well.

## Abbreviations

BLINK: Bayesian-information and linkage-disequilibrium iteratively nested keyway;
BLUE: best linear unbiased estimator; BLUP, best linear unbiased predictor;
FarmCPU: fixed and random model circulating probability unification
GAPIT: genomic association and prediction integrated tool;
GBLUP: genomic best linear unbiased predictor;
GPA: genomic prediction ability; GS, genomic selection;
GWAS: genome-wide association study;
HTP: high-throughput phenotyping;
MAS: marker assisted selection;
MIP: marginal inclusion probability;
ML: machine learning;
MTA: marker trait association;
PI: plant introduction;
PLSR: partial least squares regression;
QTL: quantitative trait locus;
RF: random forest;
SVEN: selection of variables with embedded screening using Bayesian methods;
TASSEL: trait analysis by association, evolution, and linkage;
XGBoost: extreme gradient boosting;

## ACKNOWLEDGMENTS

We thank members of the AKS group at ISU for their technical support with the field experimentation. We thank all the graduate and undergraduate students for their countless hours with phenotyping. We thank Arthur Killam for his expertise and advice in running the NIR samples. We thank Dr. Marianna Chiozza for her initial teaching on running random forests models.

## DATA AVAILABILITY STATEMENT

Data and scripts will be deposited online upon publication of the manuscript.

## AUTHOR CONTRIBUTIONS

KP and AKS conceptualized the original research, and LV and AKS conceptualized the seed protein and oil portion of the research and research questions addressed in this work. KP collected and pre-processed the data. LV and MS carried out the data analysis. SS and BG contributed to PP analysis, interpretation, and paper editing. AL contributed to seed oil and protein data collection, and GP interpretation, and paper editing. SP and HPT participated in result interpretation, paper writing and editing, and. LV wrote the first draft with feedback from AKS. All authors contributed to the paper’s development.

## SUPPLEMENTAL MATERIAL

Supplemental materials include three tables:

Table S1. Vegetation indices (VIs) computed from canopy hyperspectral reflectance.

Table S2. The complete list of all significant SNPs detected by one of the four GWAS methods, and their position in the genome.

Table S3. Distribution of significant SNPs from across the four GWAS methodologies and divided by environment and trait.

Table S4. The top 5% (13) accessions for each trait, with the maturity group information. Selections were made only from the 269 landraces that were screened in the study. For selection purposes, data for all three traits are presented. Data presented are BLUEs from across six environments (seed yield) and four environments (seed protein and oil)

## REFERENCES

Adak, A., S.C. Murray, and S.L. Anderson. 2023. Temporal phenomic predictions from unoccupied aerial systems can outperform genomic predictions. G3 13(1). doi: 10.1093/g3journal/jkac294.

Assefa, Y., N. Bajjalieh, S. Archontoulis, S. Casteel, D. Davidson, et al. 2018. Spatial Characterization of Soybean Yield and Quality (Amino Acids, Oil, and Protein) for United States. Sci. Rep. 8(1): 14653.

Ayalew, H., W. Schapaugh, T. Vuong, and H.T. Nguyen. 2022. Genome-wide association analysis identified consistent QTL for seed yield in a soybean diversity panel tested across multiple environments. Plant Genome 15(4): e20268.

Bandillo, N.B., D. Jarquin, L.G. Posadas, A.J. Lorenz, and G.L. Graef. 2023. Genomic selection performs as effectively as phenotypic selection for increasing seed yield in soybean. Plant Genome 16(1): e20285.

Bandillo, N., D. Jarquin, Q. Song, R. Nelson, P. Cregan, et al. 2015. A population structure and genome-wide association analysis on the USDA soybean germplasm collection. Plant Genome 8(3): eplantgenome2015.04.0024.

Beche, E., J.D. Gillman, Q. Song, R. Nelson, T. Beissinger, et al. 2021. Genomic prediction using training population design in interspecific soybean populations. Mol. Breed. 41(2): 15.

Bernardo, R. 2008. Molecular markers and selection for complex traits in plants: Learning from the last 20 years. Crop Sci. 48(5): 1649–1664.

Bhat, J.A., S. Ali, R.K. Salgotra, Z.A. Mir, S. Dutta, et al. 2016. Genomic Selection in the Era of Next Generation Sequencing for Complex Traits in Plant Breeding. Front. Genet. 7. doi: 10.3389/fgene.2016.00221.

Bradbury, P.J., Z. Zhang, D.E. Kroon, T.M. Casstevens, Y. Ramdoss, et al. 2007. TASSEL: software for association mapping of complex traits in diverse samples. Bioinformatics 23(19): 2633–2635.

Carley, C.N., M.J. Zubrod, S. Dutta, and A.K. Singh. 2023. Using machine learning enabled phenotyping to characterize nodulation in three early vegetative stages in soybean. Crop Sci. 63(1): 204–226.

Carroll, M.E., L.G. Riera, B.A. Miller, P.M. Dixon, B. Ganapathysubramanian, et al. 2024. Leveraging soil mapping and machine learning to improve spatial adjustments in plant breeding trials. bioRxiv: 2024.01. 03.574114. doi: 10.1101/2024.01.03.574114.

Carter, T.E., Jr, R.L. Nelson, C.H. Sneller, and Z. Cui. 2004. Genetic Diversity in Soybean. In: Shibles, R.M., Harper, J.E., Wilson, R.F., and Shoemaker, R.C., editors, Soybeans: Improvement, Production, and Uses. American Society of Agronomy, Crop Science Society of America, and Soil Science Society of America, Madison, WI, USA. p. 303–416

Chappelle, E.W., M.S. Kim, and J.E. McMurtrey. 1992. Ratio analysis of reflectance spectra (RARS): An algorithm for the remote estimation of the concentrations of chlorophyll A, chlorophyll B, and carotenoids in soybean leaves. Remote Sens. Environ. 39(3): 239– 247.

Chiozza, M.V., K.A. Parmley, R.H. Higgins, A.K. Singh, and F.E. Miguez. 2021. Comparative prediction accuracy of hyperspectral bands for different soybean crop variables: From leaf area to seed composition. Field Crops Res. 271: 108260.

Chiteri, K.O., S. Chiranjeevi, and T.Z. Jubery. 2023. Dissecting the genetic architecture of leaf morphology traits in mungbean (Vigna radiata (L.) Wizcek) using genome-wide association study. The Plant Phenome. https://acsess.onlinelibrary.wiley.com/doi/abs/10.1002/ppj2.20062.

Christenson, B.S., W.T. Schapaugh Jr, N. An, K.P. Price, V. Prasad, et al. 2016. Predicting soybean relative maturity and seed yield using canopy reflectance. Crop Sci. 56(2): 625– 643.

Clevinger, E.M., R. Biyashev, D. Haak, Q. Song, G. Pilot, et al. 2023. Identification of quantitative trait loci controlling soybean seed protein and oil content. PLoS One 18(6): e0286329.

Crain, J., S. Mondal, J. Rutkoski, R.P. Singh, and J. Poland. 2018. Combining high-throughput phenotyping and genomic information to increase prediction and selection accuracy in wheat breeding. Plant Genome 11(1): 170043.

Crossa, J., P. Pérez-Rodríguez, J. Cuevas, O. Montesinos-López, D. Jarquín, et al. 2017. Genomic Selection in Plant Breeding: Methods, Models, and Perspectives. Trends Plant Sci. 22(11): 961–975.

Dallinger, H.G., F. Löschenberger, H. Bistrich, C. Ametz, H. Hetzendorfer, et al. 2023. Predictor bias in genomic and phenomic selection. Theor. Appl. Genet. 136(11): 235.

Danilevicz, M.F., M. Gill, R. Anderson, J. Batley, M. Bennamoun, et al. 2022. Plant Genotype to Phenotype Prediction Using Machine Learning. Front. Genet. 13: 822173.

DeSalvio, A.J., A. Adak, S.C. Murray, D. Jarquín, N.D. Winans, et al. 2024. Near-infrared reflectance spectroscopy phenomic prediction can perform similarly to genomic prediction of maize agronomic traits across environments. Plant Genome: e20454.

Dobbels, A.A., and A.J. Lorenz. 2019. Soybean iron deficiency chlorosis high throughput phenotyping using an unmanned aircraft system. Plant Methods 15: 97.

Đorđević, V., M. Ćeran, J. Miladinović, S. Balešević-Tubić, K. Petrović, et al. 2019. Exploring the performance of genomic prediction models for soybean yield using different validation approaches. Mol. Breed. 39(5): 74.

Dos Santos, J.P.R., S.B. Fernandes, S. McCoy, R. Lozano, P.J. Brown, et al. 2020. Novel Bayesian networks for genomic prediction of developmental traits in biomass sorghum. G3 (Bethesda) 10(2): 769–781.

Duhnen, A., A. Gras, S. Teyssèdre, M. Romestant, B. Claustres, et al. 2017. Genomic selection for yield and seed protein content in soybean: A study of breeding program data and assessment of prediction accuracy. Crop Sci. 57(3): 1325–1337.

Durrett, T.P., C. Benning, and J. Ohlrogge. 2008. Plant triacylglycerols as feedstocks for the production of biofuels. Plant J. 54(4): 593–607.

Egli, D.B. 2023. Expanding the availability of soybean oil for renewable diesel with a high oil– low protein cultivar. Crop Sci. doi: 10.1002/csc2.21133.

Endelman, J.B. 2011. Ridge regression and other kernels for genomic selection with R package rrBLUP. Plant Genome 4(3): 250–255.

Fehr, W.R., C.E. Caviness, D.T. Burmood, and J.S. Pennington. 1971. Stage of Development Descriptions for Soybeans, *Glycine Max* (L.) Merrill ^1^. Crop Science 11(6): 929–931.

Furbank, R.T., and M. Tester. 2011. Phenomics – technologies to relieve the phenotyping bottleneck. Trends Plant Sci. 16(12): 635–644.

Gamon, J.A., L. Serrano, and J.S. Surfus. 1997. The photochemical reflectance index: an optical indicator of photosynthetic radiation use efficiency across species, functional types, and nutrient levels. Oecologia 112(4): 492–501.

Grinberg, N.F., O.I. Orhobor, and R.D. King. 2020. An evaluation of machine-learning for predicting phenotype: studies in yeast, rice, and wheat. Mach. Learn. 109(2): 251–277.

Guo, W., M.E. Carroll, A. Singh, T.L. Swetnam, N. Merchant, et al. 2021. UAS-Based Plant Phenotyping for Research and Breeding Applications. Plant Phenomics 2021: 9840192.

Hatton, N., A. Sharda, W. Schapaugh, and D. van der Merwe. 2020. Remote thermal infrared imaging for rapid screening of sudden death syndrome in soybean. Comput. Electron. Agric. 178(105738): 105738.

Hernandez, C.M., A. Correndo, P. Kyveryga, A. Prestholt, and I.A. Ciampitti. 2023. On-farm soybean seed protein and oil prediction using satellite data. Comput. Electron. Agric. 212: 108096.

Herr, A.W., A. Adak, M.E. Carroll, D. Elango, and S. Kar. 2023. Unoccupied aerial systems imagery for phenotyping in cotton, maize, soybean, and wheat breeding. Crop. https://acsess.onlinelibrary.wiley.com/doi/abs/10.1002/csc2.21028.

Herrero-Huerta, M., P. Rodriguez-Gonzalvez, and K.M. Rainey. 2020. Yield prediction by machine learning from UAS-based multi-sensor data fusion in soybean. Plant Methods. doi: 10.1186/s13007-020-00620-6;

Howard, R., and D. Jarquin. 2019. Genomic Prediction Using Canopy Coverage Image and Genotypic Information in Soybean via a Hybrid Model. Evol. Bioinform. Online 15: 1176934319840026.

Huang, M., X. Liu, Y. Zhou, R.M. Summers, and Z. Zhang. 2019. BLINK: a package for the next level of genome-wide association studies with both individuals and markers in the millions. Gigascience 8(2). doi: 10.1093/gigascience/giy154.

Hwang, E.-Y., Q. Song, G. Jia, J.E. Specht, D.L. Hyten, et al. 2014. A genome-wide association study of seed protein and oil content in soybean. BMC Genomics 15: 1.

Ibáñez, M.A., C. de Blas, L. Cámara, and G.G. Mateos. 2020. Chemical composition, protein quality and nutritive value of commercial soybean meals produced from beans from different countries: A meta-analytical study. Anim. Feed Sci. Technol. 267: 114531.

Jarquín, D., K. Kocak, L. Posadas, K. Hyma, J. Jedlicka, et al. 2014. Genotyping by sequencing for genomic prediction in a soybean breeding population. BMC Genomics 15(1): 740.

Jarquin, D., J. Specht, and A. Lorenz. 2016. Prospects of Genomic Prediction in the USDA Soybean Germplasm Collection: Historical Data Creates Robust Models for Enhancing Selection of Accessions. G3 6(8): 2329–2341.

Jin, H., X. Yang, H. Zhao, X. Song, Y.D. Tsvetkov, et al. 2023. Genetic analysis of protein content and oil content in soybean by genome-wide association study. Front. Plant Sci. 14: 1182771.

Jones, S.E., T. Ayanlade, B. Fallen, T.Z. Jubery, A. Singh, et al. 2024. Multi-Sensor and Multi-temporal High-Throughput Phenotyping for Monitoring and Early Detection of Water-Limiting Stress in Soybean. arXiv [cs.LG]. doi: 10.48550/arXiv.2402.18751.

Krause, M.D., H.-P. Piepho, K.O.G. Dias, A.K. Singh, and W.D. Beavis. 2023. Models to estimate genetic gain of soybean seed yield from annual multi-environment field trials. Züchter Genet. Breed. Res. 136(12): 252.

Kruijer, W., W. a. C.F.I.W.C.D.C. by Padraic Flood, and R. Kooke. 2023. heritability: Marker-Based Estimation of Heritability Using Individual Plant or Plot Data. https://CRAN.R-project.org/package=heritability.

Kuhn, M. 2008. Building Predictive Models in R Using the caret Package. J. Stat. Softw. 28: 1– 26.

Kusmec, A., and P.S. Schnable. 2018. FarmCPUpp: Efficient large-scale genomewide association studies. Plant Direct 2(4): e00053.

Li, S., Y. Cao, C. Wang, C. Yan, X. Sun, et al. 2023a. Genome-wide association mapping for yield-related traits in soybean (Glycine max) under well-watered and drought-stressed conditions. Front. Plant Sci. 14: 1265574.

Li, D., S. Dutta, and V. Roy. 2023b. Model Based Screening Embedded Bayesian Variable Selection for Ultra-high Dimensional Settings. J. Comput. Graph. Stat. 32(1): 61–73.

Lozano-Isla, F. 2023. inti: Tools and Statistical Procedures in Plant Science. https://CRAN.R-project.org/package=inti.

Lund, R.E. 1975. Tables for An Approximate Test for Outliers in Linear Models. Technometrics 17(4): 473–476.

Matei, G., L.G. Woyann, A.S. Milioli, I. de Bem Oliveira, A.D. Zdziarski, et al. 2018. Genomic selection in soybean: accuracy and time gain in relation to phenotypic selection. Mol. Breed. 38(9): 117.

McGowan, M., J. Wang, H. Dong, X. Liu, and Y. Jia. 2021. Plant Breeding Reviews (I. Goldman, editor). 1st ed. Wiley.

Messina, C.D., F. Technow, T. Tang, R. Totir, C. Gho, et al. 2018. Leveraging biological insight and environmental variation to improve phenotypic prediction: Integrating crop growth models (CGM) with whole genome prediction (WGP). Eur. J. Agron. 100: 151–162.

Meuwissen, T.H., B.J. Hayes, and M.E. Goddard. 2001. Prediction of total genetic value using genome-wide dense marker maps. Genetics 157(4): 1819–1829.

Miller, M.J., Q. Song, and Z. Li. 2023. Genomic selection of soybean (Glycine max) for genetic improvement of yield and seed composition in a breeding context. Plant Genome 16(4): e20384.

Monteith, J.L. 1972. Solar radiation and productivity in tropical ecosystems. J. Appl. Ecol. 9(3): 747.

Moreira, F.F., A.A. Hearst, K.A. Cherkauer, and K.M. Rainey. 2019. Improving the efficiency of soybean breeding with high-throughput canopy phenotyping. Plant Methods 15: 139.

Moreira, F.F., H.R. Oliveira, J.J. Volenec, K.M. Rainey, and L.F. Brito. 2020. Integrating High-Throughput Phenotyping and Statistical Genomic Methods to Genetically Improve Longitudinal Traits in Crops. Front. Plant Sci. 11: 681.

Nagasubramanian, K., S. Jones, A.K. Singh, S. Sarkar, A. Singh, et al. 2019. Plant disease identification using explainable 3D deep learning on hyperspectral images. Plant Methods 15: 98.

Naik, H.S., J. Zhang, A. Lofquist, T. Assefa, S. Sarkar, et al. 2017. A real-time phenotyping framework using machine learning for plant stress severity rating in soybean. Plant Methods 13: 23.

Ni, G., S. Kipp, H. Simianer, and M. Erbe. 2017. Accuracy of genomic breeding values revisited: Assessment of two established approaches and a novel one to determine the accuracy in two-step genomic prediction. J. Anim. Breed. Genet. 134(3): 242–255.

Parmley, K., K. Nagasubramanian, S. Sarkar, B. Ganapathysubramanian, and A.K. Singh. 2019. Development of Optimized Phenomic Predictors for Efficient Plant Breeding Decisions Using Phenomic-Assisted Selection in Soybean. Plant Phenomics 2019. doi: 10.34133/2019/5809404.

Peirone, L.S., G.A. Pereyra Irujo, A. Bolton, I. Erreguerena, and L.A.N. Aguirrezábal. 2018. Assessing the Efficiency of Phenotyping Early Traits in a Greenhouse Automated Platform for Predicting Drought Tolerance of Soybean in the Field. Front. Plant Sci. 9: 587.

Qin, J., F. Wang, Q. Zhao, A. Shi, T. Zhao, et al. 2022. Identification of Candidate Genes and Genomic Selection for Seed Protein in Soybean Breeding Pipeline. Front. Plant Sci. 13: 882732.

Rairdin, A., F. Fotouhi, J. Zhang, D.S. Mueller, B. Ganapathysubramanian, et al. 2022. Deep learning-based phenotyping for genome wide association studies of sudden death syndrome in soybean. Front. Plant Sci. 13: 966244.

Ravelombola, W., J. Qin, A. Shi, Q. Song, J. Yuan, et al. 2021. Genome-wide association study and genomic selection for yield and related traits in soybean. PLoS One 16(8): e0255761.

Reynolds, M., S. Chapman, L. Crespo-Herrera, G. Molero, S. Mondal, et al. 2020. Breeder friendly phenotyping. Plant Sci. 295: 110396.

Robert, P., J. Auzanneau, E. Goudemand, F.-X. Oury, B. Rolland, et al. 2022. Phenomic selection in wheat breeding: identification and optimisation of factors influencing prediction accuracy and comparison to genomic selection. Theor. Appl. Genet. 135(3): 895–914.

Rotundo, J.L., L. Borrás, M.E. Westgate, and J.H. Orf. 2009. Relationship between assimilate supply per seed during seed filling and soybean seed composition. Field Crops Res. 112(1): 90–96.

Roujean, J.-L., and F.-M. Breon. 1995. Estimating PAR absorbed by vegetation from bidirectional reflectance measurements. Remote Sens. Environ. 51(3): 375–384.

Sandhu, K.S., P.D. Mihalyov, M.J. Lewien, M.O. Pumphrey, and A.H. Carter. 2021. Combining Genomic and Phenomic Information for Predicting Grain Protein Content and Grain Yield in Spring Wheat. Front. Plant Sci. 12: 613300.

Sarkar, S., B. Ganapathysubramanian, A. Singh, F. Fotouhi, S. Kar, et al. 2024. Cyber-agricultural systems for crop breeding and sustainable production. Trends Plant Sci. 29(2): 130–149.

Shook, J., T. Gangopadhyay, L. Wu, B. Ganapathysubramanian, S. Sarkar, et al. 2021a. Crop yield prediction integrating genotype and weather variables using deep learning. PLoS One 16(6): e0252402.

Shook, J.M., D. Lourenco, and A.K. Singh. 2021b. PATRIOT: A pipeline for tracing identity-by-descent for chromosome segments to improve genomic prediction in self-pollinating crop species. Front. Plant Sci. 12: 676269.

Singh, A.K., B. Ganapathysubramanian, S. Sarkar, and A. Singh. 2018. Deep Learning for Plant Stress Phenotyping: Trends and Future Perspectives. Trends Plant Sci. 23(10): 883–898.

Singh, A., B. Ganapathysubramanian, A.K. Singh, and S. Sarkar. 2016. Machine Learning for High-Throughput Stress Phenotyping in Plants. Trends Plant Sci. 21(2): 110–124.

Singh, P., R. Kumar, S.N. Sabapathy, and A.S. Bawa. 2008. Functional and edible uses of soy protein products. Compr. Rev. Food Sci. Food Saf. 7(1): 14–28.

Singh, A.K., A. Singh, and S. Sarkar. 2021a. High-Throughput Phenotyping in Soybean. High-Throughput Crop. https://link.springer.com/chapter/10.1007/978-3-030-73734-4_7.

Singh, D.P., A.K. Singh, and A. Singh. 2021b. Plant Breeding and Cultivar Development. Elsevier Science.

Song, Q., D.L. Hyten, G. Jia, C.V. Quigley, E.W. Fickus, et al. 2015. Fingerprinting Soybean Germplasm and Its Utility in Genomic Research. G3 5(10): 1999–2006.

Song, H., X. Wang, Y. Guo, and X. Ding. 2022. G × EBLUP: A novel method for exploring genotype by environment interactions and genomic prediction. Front. Genet. 13: 972557.

Stewart-Brown, B.B., Q. Song, J.N. Vaughn, and Z. Li. 2019. Genomic Selection for Yield and Seed Composition Traits Within an Applied Soybean Breeding Program. G3 9(7): 2253–2265.

Sun, B., R. Guo, Z. Liu, X. Shi, Q. Yang, et al. 2022. Genetic variation and marker−trait association affect the genomic selection prediction accuracy of soybean protein and oil content. Front. Plant Sci. 13. doi: 10.3389/fpls.2022.1064623.

Technow, F., C.D. Messina, L.R. Totir, and M. Cooper. 2015. Integrating Crop Growth Models with Whole Genome Prediction through Approximate Bayesian Computation. PLoS One 10(6): e0130855.

Torsoni, G.B., L.E. de Oliveira Aparecido, G.M. dos Santos, A.G. Chiquitto, J.R. da Silva Cabral Moraes, et al. 2023. Soybean yield prediction by machine learning and climate. Theor. Appl. Climatol. 151(3): 1709–1725.

Vieira, C.C., R. Persa, P. Chen, and D. Jarquin. 2022. Incorporation of soil-derived covariates in progeny testing and line selection to enhance genomic prediction accuracy in soybean breeding. Front. Genet. 13: 905824.

Vogelmann, J.E., B.N. Rock, and D.M. Moss. 1993. Red edge spectral measurements from sugar maple leaves. Int. J. Remote Sens. 14(8): 1563–1575.

Vollmann, J., H. Walter, T. Sato, and P. Schweiger. 2011. Digital image analysis and chlorophyll metering for phenotyping the effects of nodulation in soybean. Comput. Electron. Agric. 75(1): 190–195.

Wang, J., and Z. Zhang. 2021. GAPIT Version 3: Boosting Power and Accuracy for Genomic Association and Prediction. Genomics Proteomics Bioinformatics 19(4): 629–640.

Wartha, C.A., and A.J. Lorenz. 2021. Implementation of genomic selection in public-sector plant breeding programs: Current status and opportunities. Crop Breed. Appl. Biotechnol. 21(spe). doi: 10.1590/1984-70332021v21sa28.

Wolc, A., and J.C.M. Dekkers. 2022. Application of Bayesian genomic prediction methods to genome-wide association analyses. Genet. Sel. Evol. 54(1): 31.

Yanarella, C.F., L. Fattel, and C.J. Lawrence-Dill. 2024. Genome-wide association studies from spoken phenotypic descriptions: a proof of concept from maize field studies. G3: jkae161.

Yang, W., H. Feng, X. Zhang, J. Zhang, J.H. Doonan, et al. 2020. Crop Phenomics and High-Throughput Phenotyping: Past Decades, Current Challenges, and Future Perspectives. Mol. Plant 13(2): 187–214.

Yang, J., J. Zeng, M.E. Goddard, N.R. Wray, and P.M. Visscher. 2017. Concepts, estimation and interpretation of SNP-based heritability. Nat. Genet. 49(9): 1304–1310.

Young, T.J., S. Chiranjeevi, D. Elango, S. Sarkar, A.K. Singh, et al. 2024. Soybean canopy stress classification using 3D point cloud data. Agronomy (Basel) 14(6): 1181.

Young, T.J., T.Z. Jubery, C.N. Carley, M. Carroll, S. Sarkar, et al. 2023. “Canopy fingerprints” for characterizing three-dimensional point cloud data of soybean canopies. Front. Plant Sci. 14: 1141153.

Zhang, H., W. Goettel, Q. Song, H. Jiang, Z. Hu, et al. 2020. Selection of GmSWEET39 for oil and protein improvement in soybean. PLoS Genet. 16(11): e1009114.

Zhang, S., D. Hao, S. Zhang, D. Zhang, H. Wang, et al. 2021. Genome-wide association mapping for protein, oil and water-soluble protein contents in soybean. Mol. Genet. Genomics 296(1): 91–102.

Zhang, J., Q. Song, P.B. Cregan, and G.-L. Jiang. 2016. Genome-wide association study, genomic prediction and marker-assisted selection for seed weight in soybean (Glycine max). Theor. Appl. Genet. 129(1): 117–130.

Zhang, J., X. Wang, Y. Lu, S.J. Bhusal, Q. Song, et al. 2018. Genome-wide Scan for Seed Composition Provides Insights into Soybean Quality Improvement and the Impacts of Domestication and Breeding. Mol. Plant 11(3): 460–472.

Zhang, T., T. Wu, L. Wang, B. Jiang, C. Zhen, et al. 2019. A Combined Linkage and GWAS Analysis Identifies QTLs Linked to Soybean Seed Protein and Oil Content. Int. J. Mol. Sci. 20(23). doi: 10.3390/ijms20235915.

Zhou, J., H. Mou, J. Zhou, M.L. Ali, H. Ye, et al. 2021. Qualification of Soybean Responses to Flooding Stress Using UAV-Based Imagery and Deep Learning. Plant Phenomics 2021: 9892570.

Zhou, J., J. Zhou, H. Ye, M.L. Ali, H.T. Nguyen, et al. 2020. Classification of soybean leaf wilting due to drought stress using UAV-based imagery. Comput. Electron. Agric. 175: 105576.

Zhu, X., W.L. Leiser, V. Hahn, and T. Würschum. 2021. Phenomic selection is competitive with genomic selection for breeding of complex traits. The Plant Phenome Journal 4(1). doi: 10.1002/ppj2.20027.

