## Supplemental Tables for "Genomic and Phenomic Prediction for Soybean Seed Yield, Protein, and Oil"

**Table S1. Vegetation indices (VI) computed from canopy hyperspectral reflectance (R).**

| Vegetation Index | Formula[a] |
|---|---|
| Normalized Vegetation Index (NDVI) | $R_{780}$-$R_{680}$/$R_{780}$+$R_{680}$ |
| Normalized Water Index (NWI) | $R_{970}$-$R_{900}$/$R_{970}$+$R_{900}$ |
| Photochemical Reflectance Index (PRI) | $R_{531}$-$R_{570}$/$R_{531}$+$R_{570}$ |
| Ratio Analysis of Reflectance Spectra Chlorophyll a (RARSa) | $R_{675}$ /$R_{700}$ |
| Ratio Analysis of Reflectance Spectra Chlorophyll b (RARSb) | $R_{675}$ /($R_{650}$*$R_{700}$) |
| Ratio Analysis of Reflectance Spectra Chlorophyll c (RARSb) | $R_{760}$/$R_{500}$ |
| Vogelmann Red Edge Index 2 (VREI2) | $R_{734}$-$R_{747}$/$R_{715}$+$R_{726}$ |
| Normalized Lignin Index (NDLI) | [log(1/$R_{1754}$)-log(1/$R_{1680}$)]/ [log(1/$R_{1754}$)+log(1/$R_{1680}$)] |
| Normalized Multi-band Drought Index (NMDI) | [$R_{860}$-($R_{1640}$ – $R_{2130}$)]/[$R_{860}$+($R_{1640}$– $R_{2130}$)] |
| Red Normalized Difference Vegetation Index (RDVI) | $R_{800}$-$R_{670}$/$R_{800}$+$R_{670}$ |

[a]R indicates the wavelength of the spectral reflectance

**Table S2.** List of all significant Marker Trait Associations (MTAs) reported based on LOD greater than 5.87 or an MIP greater than 0.5 across all four programs.

| MTA | Chromosome | Position | Trait | Environment | MIP/LOD | GWAS Method |
|---|---|---|---|---|---|---|
| ss715578520 | 1 | 1475349 | Oil | Agron17 | 0.9862 | SVEN |
| ss715578750 | 1 | 2364655 | Oil | All | 6.4622 | BLINK |
| ss715580339 | 1 | 53105494 | Oil | Lewis17 | 6.6251 | FarmCPU |
| ss715581369 | 2 | 16263148 | Oil | All | 8.0960 | BLINK |
| ss715582619 | 2 | 40617738 | Oil | Lewis17 | 6.6850 | BLINK |
| ss715585001 | 3 | 26454112 | Yield | Lewis16 | 8.2562 | FarmCPU |
| ss715585985 | 3 | 38887540 | Yield | Agron16 | 7.1367 | FarmCPU |
| | | | | | 7.7594 | BLINK |
| ss715589731 | 4 | 1013942 | Protein | All | 9.3127 | FarmCPU |
| ss715587027 | 4 | 1040604 | Protein | All | 6.9640 | BLINK |
| ss715589701 | 4 | 9660184 | Oil | All | 5.9325 | BLINK |
| | | | | | 0.8985 | SVEN |
| | | | | Agron17 | 12.1208 | FarmCPU |
| | | | | | 13.2240 | BLINK |
| | | | | | 0.5117 | SVEN |
| ss715589702 | 4 | 9674684 | Oil | Lewis16 | 0.9980 | SVEN |
| ss715587189 | 4 | 16536652 | Yield | All | 8.7933 | FarmCPU |
| ss715587823 | 4 | 39484148 | Yield | Mars17 | 9.1797 | BLINK |
| ss715587893 | 4 | 41351742 | Yield | Lewis16 | 9.9950 | FarmCPU |
| | | | | | 6.0383 | BLINK |
| | | | | | 0.8974 | SVEN |
| ss715588739 | 4 | 49650372 | Oil | All | 7.2980 | FarmCPU |
| ss715588918 | 4 | 51108220 | Oil | All | 0.6045 | SVEN |
| ss715592605 | 5 | 213770 | Protein | All | 7.4795 | FarmCPU |
| ss715592443 | 5 | 2372200 | Protein | All | 6.9082 | FarmCPU |
| ss715589830 | 5 | 2849405 | Oil | Lewis16 | 6.0121 | TASSEL |
| ss715590626 | 5 | 31450644 | Yield | Lewis16 | 7.2719 | BLINK |
| ss715591274 | 5 | 36150537 | Protein | Lewis16 | 6.8145 | FarmCPU |
| | | | | | 0.9091 | SVEN |
| ss715591885 | 5 | 40327284 | Oil | Cas17 | 6.3590 | FarmCPU |
| ss715591862 | 5 | 40519175 | Yield | All | 7.7196 | FarmCPU |
| ss715591697 | 5 | 41415712 | Oil | All | 0.8246 | SVEN |
| | | | | Agron17 | 8.2289 | FarmCPU |
| | | | | | 7.2058 | BLINK |
| | | | | | 0.9963 | SVEN |
| ss715592683 | 6 | 10250655 | Oil | Lewis16 | 7.0618 | FarmCPU |
| ss715594386 | 6 | 41376571 | Yield | Cas17 | 6.8346 | FarmCPU |
| ss715596765 | 7 | 20339164 | Yield | Agron16 | 8.3020 | FarmCPU |
| ss715596766 | 7 | 20392261 | Yield | Lewis17 | 7.0341 | TASSEL |
| ss715599603 | 8 | 1475148 | Yield | Cas17 | 6.1172 | FarmCPU |
| ss715600736 | 8 | 2180765 | Yield | Lewis17 | 6.0417 | TASSEL |

| | | | | | | |
|---|---|---|---|---|---|---|
| ss715602559 | 8 | 5088534 | Yield | Cas17 | 7.7333 | BLINK |
| ss715602691 | 8 | 7574121 | Oil | Lewis17 | 0.6075 | SVEN |
| ss715602751 | 8 | 8284178 | Oil | Cas17 | 7.3140 | FarmCPU |
| ss715602786 | 8 | 8657875 | Oil | Lewis16 | 0.5735 | SVEN |
| ss715602805 | 8 | 8847325 | Oil | All | 0.6315 | SVEN |
| | | | | Lewis16 | 7.3713 | BLINK |
| ss715599801 | 8 | 16261932 | Yield | Lewis17 | 0.8453 | SVEN |
| ss715600183 | 8 | 19241199 | Oil | Cas17 | 6.3362 | FarmCPU |
| ss715600189 | 8 | 19249288 | Protein | Lewis17 | 6.7097 | BLINK |
| ss715600934 | 8 | 22636972 | Oil | Cas17 | 6.7087 | BLINK |
| ss715602480 | 8 | 47076199 | Yield | Lewis16 | 7.7679 | BLINK |
| ss715605125 | 9 | 5171096 | Oil | Agron17 | 5.9683 | FarmCPU |
| ss715605308 | 9 | 6517142 | Protein | All | 0.7435 | SVEN |
| ss715605513 | 10 | 1072928 | Protein | Lewis17 | 6.2887 | FarmCPU |
| ss715605712 | 10 | 1666457 | Yield | Agron16 | 0.5324 | SVEN |
| | | | | Cas17 | 6.6673 | BLINK |
| ss715606401 | 10 | 37188800 | Protein | Lewis16 | 8.3136 | FarmCPU |
| ss715607389 | 10 | 44507098 | Oil | Lewis17 | 0.7139 | SVEN |
| ss715609637 | 11 | 2124435 | Yield | Mars17 | 6.4195 | BLINK |
| | | | | | 0.5491 | SVEN |
| ss715610619 | 11 | 3898977 | Yield | Mars17 | 6.2048 | TASSEL |
| ss715611003 | 11 | 6565408 | Yield | Cas17 | 0.6117 | SVEN |
| ss715611078 | 11 | 7272601 | Yield | Mars17 | 5.9501 | BLINK |
| ss715611224 | 11 | 8293886 | Oil | Lewis16 | 8.8113 | BLINK |
| ss715611247 | 11 | 8627594 | Yield | Agron16 | 6.3473 | BLINK |
| ss715611267 | 11 | 8876028 | Oil | Cas17 | 7.3495 | FarmCPU |
| ss715611302 | 11 | 9421766 | Yield | Lewis17 | 0.8718 | SVEN |
| ss715610552 | 11 | 34172468 | Oil | Agron17 | 6.3931 | BLINK |
| ss715612413 | 12 | 34373853 | Yield | Cas17 | 9.5759 | BLINK |
| ss715617022 | 13 | 14987056 | Protein | All | 8.8835 | FarmCPU |
| ss715615568 | 13 | 18519484 | Protein | All | 8.7334 | FarmCPU |
| | | | | | 6.0175 | BLINK |
| | | | | | 0.9979 | SVEN |
| | | | | Lewis16 | 7.8073 | FarmCPU |
| | | | | | 0.8644 | SVEN |
| | | | | Lewis17 | 0.5319 | SVEN |
| | | | | Cas17 | 0.9926 | SVEN |
| ss715614058 | 13 | 22858156 | Yield | Lewis17 | 5.8808 | FarmCPU |
| | | | | All | 6.5733 | BLINK |
| ss715614768 | 13 | 29216173 | Yield | Cas17 | 6.6770 | BLINK |
| ss715614988 | 13 | 30502735 | Yield | Lewis17 | 6.4571 | FarmCPU |
| ss715615470 | 13 | 34057492 | Oil | All | 7.9759 | TASSEL |
| | | | | Lewis16 | 9.5897 | TASSEL |
| | | | | Lewis17 | 6.1430 | TASSEL |
| ss715615595 | 13 | 35077503 | Oil | All | 6.3384 | BLINK |

| | | | | | | |
|---|---|---|---|---|---|---|
| ss715616016 | 13 | 38165515 | Yield | All | 0.7549 | SVEN |
| ss715619757 | 14 | 640339 | Yield | Lewis16 | 0.8879 | SVEN |
| ss715617325 | 14 | 1009520 | Yield | Lewis16 | 6.8324 | FarmCPU |
| ss715620089 | 14 | 9099832 | Yield | Cas17 | 6.7569 | FarmCPU |
| ss715617374 | 14 | 10095909 | Oil | All | 6.8822 | FarmCPU |
| ss715617374 | 14 | 10095909 | Oil | All | 0.9419 | SVEN |
| ss715617374 | 14 | 10095909 | Oil | Agron17 | 0.8892 | SVEN |
| ss715617374 | 14 | 10095909 | Oil | Cas17 | 6.8485 | FarmCPU |
| ss715621777 | 15 | 3846538 | Oil | All | 8.9913 | FarmCPU |
| ss715621777 | 15 | 3846538 | Oil | All | 0.9531 | SVEN |
| ss715621777 | 15 | 3846538 | Oil | Lewis16 | 6.5498 | BLINK |
| ss715621777 | 15 | 3846538 | Oil | Agron17 | 0.9606 | SVEN |
| ss715621777 | 15 | 3846538 | Protein | Agron17 | 0.8571 | SVEN |
| ss715621781 | 15 | 3863922 | Protein | Lewis17 | 6.1451 | FarmCPU |
| ss715622562 | 15 | 4926375 | Protein | All | 7.6288 | FarmCPU |
| ss715622682 | 15 | 5014514 | Oil | Lewis17 | 0.5555 | SVEN |
| ss715622682 | 15 | 5014514 | Oil | Cas17 | 0.9261 | SVEN |
| ss715623160 | 15 | 8751332 | Oil | Lewis16 | 6.0841 | TASSEL |
| ss715620177 | 15 | 10213666 | Oil | Lewis16 | 6.9519 | FarmCPU |
| ss715620226 | 15 | 10520900 | Yield | All | 7.9014 | FarmCPU |
| ss715620226 | 15 | 10520900 | Yield | All | 6.3387 | BLINK |
| ss715620227 | 15 | 10545687 | Yield | Lewis17 | 7.3767 | FarmCPU |
| ss715620227 | 15 | 10545687 | Yield | Lewis17 | 8.2705 | BLINK |
| ss715620227 | 15 | 10545687 | Yield | Lewis17 | 0.5007 | SVEN |
| ss715620229 | 15 | 10553131 | Yield | Lewis16 | 10.3388 | FarmCPU |
| ss715620229 | 15 | 10553131 | Yield | Lewis16 | 12.2168 | BLINK |
| ss715620229 | 15 | 10553131 | Yield | Cas17 | 6.8575 | FarmCPU |
| ss715620238 | 15 | 10702955 | Oil | Lewis17 | 8.6277 | FarmCPU |
| ss715620322 | 15 | 11404032 | Yield | Lewis16 | 0.8992 | SVEN |
| ss715623414 | 16 | 305626 | Protein | Agron17 | 0.7728 | SVEN |
| ss715625179 | 16 | 509530 | Oil | All | 7.9248 | FarmCPU |
| ss715623475 | 16 | 1498023 | Oil | Agron17 | 0.5115 | SVEN |
| ss715623476 | 16 | 1500606 | Oil | All | 0.7680 | SVEN |
| ss715624371 | 16 | 3113454 | Yield | Agron17 | 0.8988 | SVEN |
| ss715623933 | 16 | 28573947 | Yield | All | 6.2158 | FarmCPU |
| ss715624165 | 16 | 29631736 | Oil | Lewis17 | 0.5421 | SVEN |
| ss715624192 | 16 | 29870849 | Yield | All | 6.6009 | FarmCPU |
| ss715624192 | 16 | 29870849 | Yield | Cas17 | 9.2232 | FarmCPU |
| ss715624192 | 16 | 29870849 | Yield | All | 7.0367 | BLINK |
| ss715624199 | 16 | 29940504 | Yield | All | 0.6544 | SVEN |
| ss715624199 | 16 | 29940504 | Yield | Lewis17 | 6.5525 | BLINK |
| ss715624199 | 16 | 29940504 | Yield | Lewis17 | 0.9376 | SVEN |
| ss715624199 | 16 | 29940504 | Yield | Cas17 | 0.9930 | SVEN |
| ss715624379 | 16 | 31181902 | Yield | Agron16 | 0.7019 | SVEN |
| ss715624562 | 16 | 32882556 | Oil | All | 6.9409 | BLINK |

| | | | | | | |
|---|---|---|---|---|---|---|
| ss715624996 | 16 | 37570986 | Oil | Agron17 | 5.8953 | FarmCPU |
| ss715626431 | 17 | 2217986 | Oil | Lewis17 | 6.6633 | BLINK |
| ss715626741 | 17 | 3244943 | Yield | Lewis17 | 0.8909 | SVEN |
| ss715626753 | 17 | 3287941 | Yield | Lewis16 | 6.2624 | TASSEL |
| | | | | | 8.8023 | FarmCPU |
| | | | | | 1.0000 | SVEN |
| ss715626757 | 17 | 3300391 | Yield | Lewis17 | 6.4416 | TASSEL |
| ss715626802 | 17 | 3433472 | Yield | Agron17 | 0.7076 | SVEN |
| ss715627800 | 17 | 3846471 | Protein | All | 6.0379 | BLINK |
| | | | | | 0.9983 | SVEN |
| | | | | Lewis16 | 6.8083 | FarmCPU |
| | | | | | 0.9916 | SVEN |
| | | | | Lewis17 | 11.1782 | BLINK |
| | | | | | 0.9876 | SVEN |
| ss715628250 | 17 | 8125412 | Oil | Cas17 | 0.8160 | SVEN |
| ss715628317 | 17 | 8672815 | Oil | Lewis17 | 6.6187 | FarmCPU |
| ss715628318 | 17 | 8676537 | Protein | All | 6.2846 | FarmCPU |
| ss715625851 | 17 | 12128041 | Oil | Lewis16 | 0.5376 | SVEN |
| ss715625889 | 17 | 12540814 | Oil | Lewis16 | 6.0707 | FarmCPU |
| ss715627506 | 17 | 38654233 | Oil | Lewis16 | 7.4527 | FarmCPU |
| ss715627546 | 17 | 38942962 | Yield | Lewis16 | 7.3265 | BLINK |
| ss715627547 | 17 | 38944808 | Yield | Lewis16 | 8.4357 | FarmCPU |
| | | | | | 0.5956 | SVEN |
| ss715630114 | 18 | 2798728 | Yield | All | 6.5869 | BLINK |
| | | | | | 0.9977 | SVEN |
| | | | | Cas17 | 7.1669 | FarmCPU |
| ss715630802 | 18 | 4813092 | Oil | All | 0.8864 | SVEN |
| ss715631416 | 18 | 48842474 | Oil | Lewis16 | 0.5024 | SVEN |
| ss715632014 | 18 | 54249904 | Yield | Agron16 | 0.5542 | SVEN |
| ss715632030 | 18 | 54334495 | Yield | All | 8.2812 | FarmCPU |
| ss715633641 | 19 | 2960221 | Yield | Lewis17 | 7.2163 | TASSEL |
| ss715633724 | 19 | 31684890 | Yield | All | 9.5644 | FarmCPU |
| ss715634225 | 19 | 35109354 | Yield | Lewis17 | 0.8318 | SVEN |
| ss715634486 | 19 | 36560503 | Yield | Lewis17 | 6.1593 | FarmCPU |
| | | | | | 8.1552 | BLINK |
| ss715634507 | 19 | 36781528 | Yield | Agron17 | 0.7114 | SVEN |
| ss715634757 | 19 | 38801914 | Oil | Cas17 | 0.5271 | SVEN |
| ss715634834 | 19 | 39563987 | Yield | Agron16 | 5.9864 | FarmCPU |
| ss715635419 | 19 | 45159526 | Protein | All | 7.5949 | FarmCPU |
| | | | | | 12.1400 | BLINK |
| | | | | Lewis16 | 11.8604 | BLINK |
| | | | | Lewis17 | 8.5134 | FarmCPU |
| | | | | | 7.3977 | BLINK |
| ss715636617 | 20 | 8318718 | Yield | Agron17 | 0.8484 | SVEN |
| ss715636741 | 20 | 12866530 | Oil | Agron17 | 0.9888 | SVEN |

| | | | | | | |
|---|---|---|---|---|---|---|
| ss715639075 | 20 | 19385331 | Oil | All | 0.6836 | SVEN |
| | | | | Lewis16 | 6.8940 | FarmCPU |
| ss715637055 | 20 | 25444475 | Oil | All | 9.1457 | FarmCPU |
| | | | | Agron17 | 9.6553 | BLINK |
| | | | | Cas17 | 8.7332 | FarmCPU |
| ss715637185 | 20 | 29719631 | Yield | Lewis16 | 6.9456 | BLINK |
| ss715637217 | 20 | 30546685 | Protein | Lewis16 | 7.3174 | FarmCPU |
| ss715637499 | 20 | 34907808 | Protein | Lewis17 | 6.3160 | FarmCPU |
| ss715638023 | 20 | 39341961 | Protein | All | 6.1464 | BLINK |
| ss715638822 | 20 | 46890422 | Oil | Lewis16 | 8.1145 | BLINK |

**Table S3. Distribution of significant MTAs across the four GWAS methodologies (SVEN, MLM-TASSEL, FarmCPU, BLINK) used to run GWAS. Totals represent the number of unique MTAs after accounting for overlapping MTAs across methods, environments, and traits.**

| Trait | Environment | SVEN | TASSEL | FarmCPU | BLINK | Total[§] |
|---|---|---|---|---|---|---|
| Yield | Combined | 3 | 0 | 7 | 4 | 11 |
| | Agron16 | 3 | 0 | 3 | 2 | 7 |
| | Lewis16 | 5 | 1 | 6 | 6 | 12 |
| | Agron17 | 4 | 0 | 0 | 0 | 4 |
| | Lewis17 | 6 | 4 | 4 | 3 | 13 |
| | Cas17 | 2 | 0 | 6 | 4 | 12 |
| | Mars17 | 1 | 1 | 0 | 3 | 4 |
| | Total[§] | 22 | 6 | 24 | 22 | 56 |
| Protein | Combined | 3 | 0 | 8 | 5 | 12 |
| | Lewis16 | 3 | 0 | 5 | 1 | 6 |
| | Agron17 | 2 | 0 | 0 | 0 | 2 |
| | Lewis17 | 2 | 0 | 4 | 3 | 7 |
| | Cas17 | 1 | 0 | 0 | 0 | 1 |
| | Total[§] | 6 | 0 | 15 | 6 | 21 |
| Oil | Combined | 9 | 1 | 5 | 5 | 17 |
| | Lewis16 | 4 | 3 | 5 | 4 | 16 |
| | Agron17 | 7 | 0 | 4 | 4 | 11 |
| | Lewis17 | 4 | 1 | 3 | 2 | 10 |
| | Cas17 | 3 | 0 | 6 | 1 | 10 |
| | Total[§] | 22 | 3 | 21 | 15 | 51 |
| | Grand Total[§] | 49 | 9 | 60 | 43 | 127 |

[§]Totals with overlapping MTAs across methods, environments, and traits removed

**Table S4.** The top 5% (13) accessions for each trait, with the maturity group information. Selections were made only from the 269 landraces that were screened in the study. For selection purposes, data for all three traits are presented. Data presented are BLUEs from across six environments (seed yield) and four environments (seed protein and oil)

| Trait | Accession | MG | Seed Yield (kg ha$^{-1}$) | Protein (% dry basis) | Oil (% dry basis) |
|---|---|---|---|---|---|
| **Top 5% Seed Yield** | PI 597482 | III | 3635.7 | 37.3 | 21.4 |
| | PI 518751 | II | 3273.2 | 39.6 | 20.5 |
| | PI 561370 | III | 3209.4 | 40.5 | 20.3 |
| | PI 253660B | III | 3039.5 | 40.1 | 19.5 |
| | PI 538400 | II | 2988.7 | 40.9 | 20.0 |
| | PI 427136 | III | 2970.9 | 41.4 | 20.7 |
| | PI 189930 | II | 2970.5 | 39.7 | 19.7 |
| | PI 574486 | III | 2964.2 | 40.9 | 20.1 |
| | PI 574480B | III | 2961.4 | 40.4 | 20.3 |
| | PI 592910 | II | 2960.9 | 40.9 | 20.1 |
| | PI 578367 | III | 2868.1 | 42.3 | 19.6 |
| | PI 578499B | II | 2838.7 | 41.1 | 19.0 |
| | PI 578416 | II | 2833.4 | 39.4 | 20.5 |
| **Top 5% Seed Protein** | PI 437377 | III | 2076.5 | 48.6 | 16.6 |
| | PI 549021A | III | 1500.1 | 47.5 | 14.8 |
| | PI 430597 | II | 468.7 | 47.2 | 14.8 |
| | PI 437145B | II | 1653.8 | 47.1 | 16.1 |
| | PI 416868A | III | 1384.0 | 46.5 | 16.5 |
| | PI 506887 | III | 1673.5 | 46.4 | 16.5 |
| | PI 391577 | II | 1076.8 | 46.3 | 14.8 |
| | PI 87618 | III | 1920.7 | 45.9 | 16.9 |
| | PI 567267A | II | 924.9 | 45.8 | 16.0 |
| | PI 417559 | III | 1816.0 | 45.8 | 15.1 |
| | PI 567538B | II | 1527.0 | 45.6 | 16.7 |
| | PI 567366B | IV | 1719.2 | 45.6 | 16.9 |
| | PI 518757 | III | 2011.4 | 45.6 | 15.9 |
| **Top 5% Seed Oil** | PI 347552B | I | 2072.7 | 37.1 | 22.7 |
| | PI 84921 | II | 2699.0 | 38.0 | 21.6 |
| | PI 68788 | II | 2524.7 | 39.5 | 21.5 |
| | PI 437592 | II | 2592.7 | 39.8 | 21.5 |
| | PI 290134 | I | 2119.8 | 38.6 | 21.5 |
| | PI 597482 | III | 3635.7 | 37.3 | 21.4 |
| | PI 89130 | III | 2426.4 | 40.1 | 21.4 |
| | PI 438139 | II | 2549.2 | 40.1 | 21.4 |
| | PI 92603 | II | 2395.4 | 38.7 | 21.3 |
| | PI 92611 | II | 2225.8 | 39.5 | 21.3 |
| | PI 70463 | II | 2446.0 | 40.2 | 21.2 |
| | PI 80461 | III | 2666.0 | 38.5 | 21.2 |
| | PI 437122 | II | 2349.3 | 40.0 | 21.1 |
